# The Logic of Thalamic Inputs onto the Molecular Taxonomy of Cortical Neurons Reveals a Visual Hierarchy

**DOI:** 10.64898/2026.06.12.731929

**Authors:** Xinxin Ge, Xing Wei, Baixi Ruan, Qin Ying Wu, Siyuan Chang, Nicole Tsai, Shaobo Zhang, Xin Duan, Massimo Scanziani

## Abstract

The hierarchical organization of sensory cortices and the rich molecular taxonomy of their cell types are defining features of the mammalian cortex. Cortical areas along this hierarchy are reciprocally connected via the thalamus through bottom-up and top-down projections. The logic through which these projections map onto the cellular taxonomy of the cortex is, however, poorly understood. Here we combine an anterograde transsynaptic tracer with spatial transcriptomics to reveal the molecular and spatial identity of mouse visual cortical neurons downstream of thalamic starter neurons across visual cortical areas. Distinct thalamic inputs target characteristic sets of molecularly defined cortical neurons, forming bottom-up or top-down “signatures” defined by cell-type composition and the ratio of GABAergic to glutamatergic neurons. These signatures reveal a hierarchy spanning thalamic nuclei and visual cortical areas, independently predicted by the molecular and cellular similarities between cortical areas. This work uncovers basic principles of how bottom-up and top-down thalamic inputs map onto the cellular taxonomy in the visual cortex and establishes a cellular framework for the cortical hierarchy.

## Main

A fundamental principle of cortical organization in the mammalian brain is the hierarchical structure of its sensory areas^1,2^. This hierarchy is defined by two streams of sensory information flowing in opposite directions: bottom-up and top-down^3^. Primary sensory cortical areas receive ascending inputs from the sensory periphery via primary thalamic nuclei and forward this information to higher cortical areas via higher-order thalamic nuclei, forming the bottom-up stream. Conversely, higher cortical areas send information back to lower areas through higher-order thalamic nuclei, constituting the top-down stream. That is, the thalamus serves as a central hub for the transmission of bottom-up and top-down signals throughout the cortical hierarchy^4,5^. The identity of cortical neurons targeted by these two streams of information remains poorly understood. Recent advances in single-cell transcriptomics have uncovered an unprecedented molecular diversity of mouse cortical neurons^6–10^.

While these discoveries illustrate the rich cellular composition of the mammalian cortex, they also highlight a major gap in our understanding of the cortical circuit architecture, namely, how the two principal streams of sensory information, the bottom-up and the top-down, map onto this newly revealed cellular taxonomy. Is there a characteristic pattern of molecularly distinct cortical neurons targeted by the bottom-up or top-down stream? And if so, is this pattern conserved across the cortical hierarchy?

We tackled these questions in the mouse visual system, where the hierarchical organization of visual cortical areas is well established^11–15^ and, importantly, where the cellular make-up of these areas has been cataloged in precisely defined molecular types^7,8,10^. The dorsolateral geniculate nucleus (dLGN) is the primary thalamic nucleus relaying bottom-up visual information originating in the retina to primary visual cortex (V1)^16,17^. The pulvinar (also known as lateral posterior (LP) in rodents) is the higher-order thalamic nucleus that links V1 and higher visual cortical areas (HVCs) along the hierarchy, via bottom-up and top-down projections^18–20^. Here, we integrate a recently developed anterograde transsynaptic tracer, a reengineered wheat germ agglutinin fused to mCherry (mWGA–mCherry hereafter)^21^, with spatial transcriptomics (MERFISH)^22^. This integrated technical platform, named “TransA-MERFISH”, enables simultaneous spatial registration and molecular classification of synaptically connected neurons downstream of regionally or genetically defined starter neurons in a high-throughput manner. Through the TransA-MERFISH platform, we not only discovered distinct patterns of cortical neurons targeted by bottom-up and top-down streams but also revealed characteristic signatures of how these two streams map onto the rich diversity of cortical neurons, irrespective of the thalamic nuclei of origin or the targeted cortical area. These results allow us to establish a hierarchy of cortical areas based on the molecular composition of neurons targeted by thalamic inputs.

### Integrating mWGA-mCherry with MERFISH to map thalamocortical connectivity

To identify visual cortical neurons that receive direct input from the dLGN or LP, we adapted an anterograde transsynaptic tracing technology, AAV-encoded mWGA-mCherry, originally established for retino-tectal projection to the thalamocortical circuit^21^. AAV-packaged mWGA-mCherry was injected into either thalamic nucleus (Fig. 1a, b; Methods). Upon expression, the fused protein is transported along axons and crosses synapses to label monosynaptically connected neurons in the anterograde direction^21^. The molecular identity of the fluorescently labeled visual cortical neurons can then be revealed using spatial transcriptomics.

**Fig. 1.**
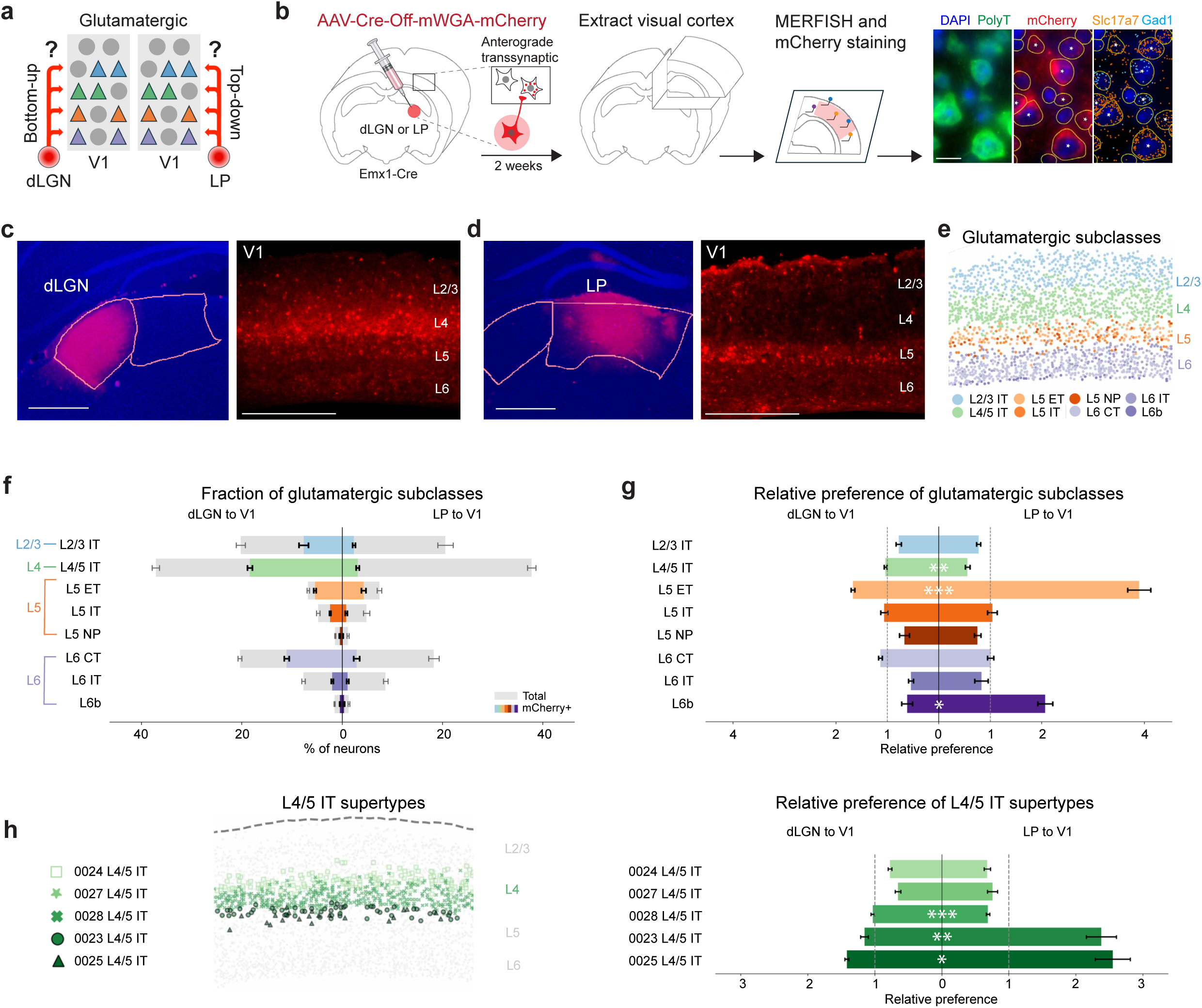
Distinct patterns of glutamatergic V1 neurons targeted by dLGN and LP. **a,** Schematic illustrating the question addressed in this figure: which molecularly defined glutamatergic neurons are targeted by bottom-up dLGN input or top-down LP input? **b,** Schematic of the TransA-MERFISH experimental approach for thalamocortical circuit mapping. Asterisks in the final panel denote cells with a posterior probability of being mCherry⁺ greater than 0.8. Scale bar, 10µm. **c, d,** Representative images showing mWGA-mCherry expression at the injection site in the thalamus (left) and anterogradely transferred mWGA-mCherry signal in V1 (right), two weeks after injection into the dLGN (**c**) or LP (**d**). Scale bars, 0.5 mm. Image intensity ranges were not matched between panels. **e,** Spatial distribution of major glutamatergic subclasses defined by their transcriptomic profile in the same example V1 section shown in **c**. IT: intratelencephalic, ET: extratelencephalic, NP: near projecting, CT: corticothalamic. See **c** for scale bar. **f,** Fractions of mCherry⁺ neurons across V1 glutamatergic subclasses following dLGN (left) or LP (right) injections. Gray bars and colored bars indicate the relative abundance of each subclass, and of the mCherry^+^ neurons, respectively, within the total glutamatergic population (right and left are different data sets). **g,** Relative preference of dLGN (left) and LP (right) inputs for each glutamatergic subclass in V1. Relative preference was calculated as the fraction of mCherry⁺ neurons in each subclass normalized to the fraction of mCherry⁺ neurons across all glutamatergic neurons. Significance was determined using two-sided Welch’s t-tests on log-transformed data, adjusted for multiple comparisons using Benjamini–Hochberg false discovery rate (FDR) correction. **h,** Left, spatial distribution of L4/5 IT supertypes in the same example V1 section shown in **c** and **e**. Note the laminar distribution of these supertypes within L4. Right, relative preference of dLGN and LP inputs for each L4/5 IT supertypes in V1. Relative preference was calculated as the fraction of mCherry⁺ neurons in each supertype normalized to the fraction of mCherry⁺ neurons across all L4/5 IT neurons. Significance was determined using two-sided Welch’s t-tests on log-transformed data, adjusted with FDR correction. dLGN to V1: n = 4, LP to V1: n = 5. Here and throughout the figure legends, “n” refers to the number of animals. Bars represent mean ± s.e.m. across animals. \**P* < 0.05, \*\**P* < 0.001, \*\*\**P* < 0.0001. Full details on statistical models and *P* values are provided in Supplementary Table 4.

To optimize and validate the AAV-encoded mWGA-mCherry anterograde transsynaptic tracing for the thalamocortical circuit, we proceeded as follows: First, because layer 5 and 6 corticothalamic neurons project from visual cortex back to the dLGN and LP, retrograde uptake of AAV-mWGA-mCherry by these axons could confound interpretation by driving mWGA-mCherry expression in cortical neurons. To eliminate this possibility, we used Cre-Off and Cre-On virogenetic strategies that restrict mWGA-mCherry expression to thalamic neurons. In the Cre-Off configuration^23^, AAV9-Cre-Off-mWGA-mCherry was injected into the dLGN or LP of Emx1-Cre mice^24^, a mouse line in which Cre is expressed in all cortical glutamatergic neurons, including corticothalamic neurons, thereby preventing their expression of mWGA-mCherry (see Methods and Extended Data Fig. 1 for validation). Unless otherwise specified, all experiments used the Cre-Off strategy. For the Cre-On strategy, we injected Cre-dependent AAV9-DIO-mWGA-mCherry into the dLGN or LP of Crh-Cre or Calb2-Cre mice^25,26^, respectively, to enable selective expression in genetically defined subsets of thalamic neurons. Importantly, expression of the Crh-Cre or Calb2-Cre in cortex is largely restricted to non-corticothalamic populations^25,26^, minimizing the likelihood of off-target cortical starter cells. The Cre-On experiments provided independent validation of the results obtained from the Cre-Off strategy.

Second, to confirm that the anterograde transfer of mWGA-mCherry from the thalamus to the visual cortex occurred between synaptically connected neurons, we performed channelrhodopsin-assisted circuit mapping experiments (CRACM)^27^ by recording from cortical neurons in acute slices while optogenetically stimulating thalamic afferents (see Methods and Extended Data Fig. 2). The probability of evoking excitatory postsynaptic currents (EPSC) was much higher in mCherry labeled neurons than unlabelled neighboring control neurons. This was the case for both pyramidal cells and genetically labeled Sst-expressing neurons, which received much smaller thalamic EPSCs compared to pyramidal cells (Extended Data Fig. 2).

Finally, mWGA-mCherry labeling remained restricted to one synapse downstream of infected thalamic neurons up to two months after injection (see Methods and Extended Data Fig. 3), much longer than the two to three weeks interval used in the experiments described below.

Following injection of AAV mWGA-mCherry in the thalamus, visual cortical sections ipsilateral to the injection sites were collected and processed for MERFISH and immunofluorescence labeling. To resolve the molecular identity of these visual cortical neurons, we selected a panel of 300 genes based on the single-cell RNA-seq (scRNA seq) data from mouse visual cortex^7,8^ (Methods; Supplementary Table 1). The panel comprises marker genes with sufficiently strong differential expression across visual cortical cell types for a classifier to provide a robust resolution of transcriptionally defined neuronal identities (Extended Data Fig. 4). Following MERFISH imaging, cells were segmented using the polyT and DAPI signals and assigned a probability of being mWGA-mCherry positive (mCherry⁺) based on immunofluorescence intensity (Methods and Fig. 1b). To assign their molecular identity, cells were mapped to the 10x Whole Mouse Brain Cell Census Network taxonomy (CCN20230722)^28^ using MapMyCell^29^ at the level of class, subclass and supertype (Methods; Fig. 1e; Extended Data Fig. 5). Molecular identity was assigned to 106,430 neurons across visual cortex (Supplementary Table 2a, b), out of which an estimated 30,521 were mCherry⁺ (n = 19 animals; see Methods; Supplementary Table 3a, b).

Thus, anterograde tracing with the current strategy reflects monosynaptic transfer from thalamic neurons to their direct synaptic targets in the visual cortex. Together with MERFISH, this integrated approach enables comprehensive molecular characterization of visual cortical neurons receiving direct thalamic input.

### TransA-MERFISH reveals distinct patterns of molecularly defined V1 neurons targeted by dLGN and LP

Because V1 receives convergent input from the dLGN and from LP, the quintessential bottom-up and top-down thalamic inputs to this visual area, we compared the patterns of glutamatergic and GABAergic neurons targeted by each of these two thalamic nuclei (Fig. 1a)

To delineate V1 we used both the extent of thalamic innervation as well as the expression of area specific genes (Fig. 1c, d; see Methods). We first quantified the fractions of mCherry⁺ neurons among major glutamatergic subclasses in V1. Overall, dLGN targeted a higher fraction of glutamatergic neurons in V1 compared to the LP (dLGN mean ± s.e.m.: 47.95 ± 1.64%, n = 4 mice; LP: 14.97 ± 1.68%, n = 5 mice). This was also true across all individual glutamatergic subclasses except for L6b (Fig. 1f).

To determine whether, despite the overall difference in the fraction of targeted neurons, dLGN and LP projections each preferentially target distinct subclasses of V1 neurons, and within these subclasses, distinct supertypes, we used the “relative preference”. The relative preference was calculated by normalizing the mCherry⁺ fraction of each subclass or supertype to the mCherry⁺ fraction across all glutamatergic neurons, or all glutamatergic neurons within a subclass, respectively (Methods). We found that dLGN and LP projections exhibited distinct preference over individual subclasses (Fig. 1g; *F_7,56_* = 20.36, *P* = 2.67 × 10^-13^, two-way ANOVA) and supertypes (Extended Data Fig. 6). Noteworthily, while the dLGN showed a higher relative preference for the L4/5 IT subclass compared to LP, within this subclass, LP favored supertypes 0023 and 0025, both located deeper in the cortex relative to other L4/5 IT supertypes, and disfavored supertype 0028, located at an intermediate depth relative to all L4/5 IT neurons (Extended Data Fig. 6). Conversely, although LP exhibited a higher relative preference for the L5 ET subclass overall, within this subclass it disfavored supertype 0093, compared to dLGN, a supertype located in superficial layer 5 (Extended Data Fig. 6).

We next examined whether dLGN and LP also target different molecularly defined GABAergic neurons in V1 (Fig. 2a). In contrast to glutamatergic neurons where each subclass shows a clear laminar distribution that can be assigned to a cortical layer, GABAergic subclasses and supertypes lack an obvious laminar organization (Fig. 2b and Extended Data Fig. 7). Therefore, each individual GABAergic neuron was assigned to a cortical layer based on the identity of the surrounding glutamatergic neurons (Fig. 2b; Methods).

**Fig. 2.**
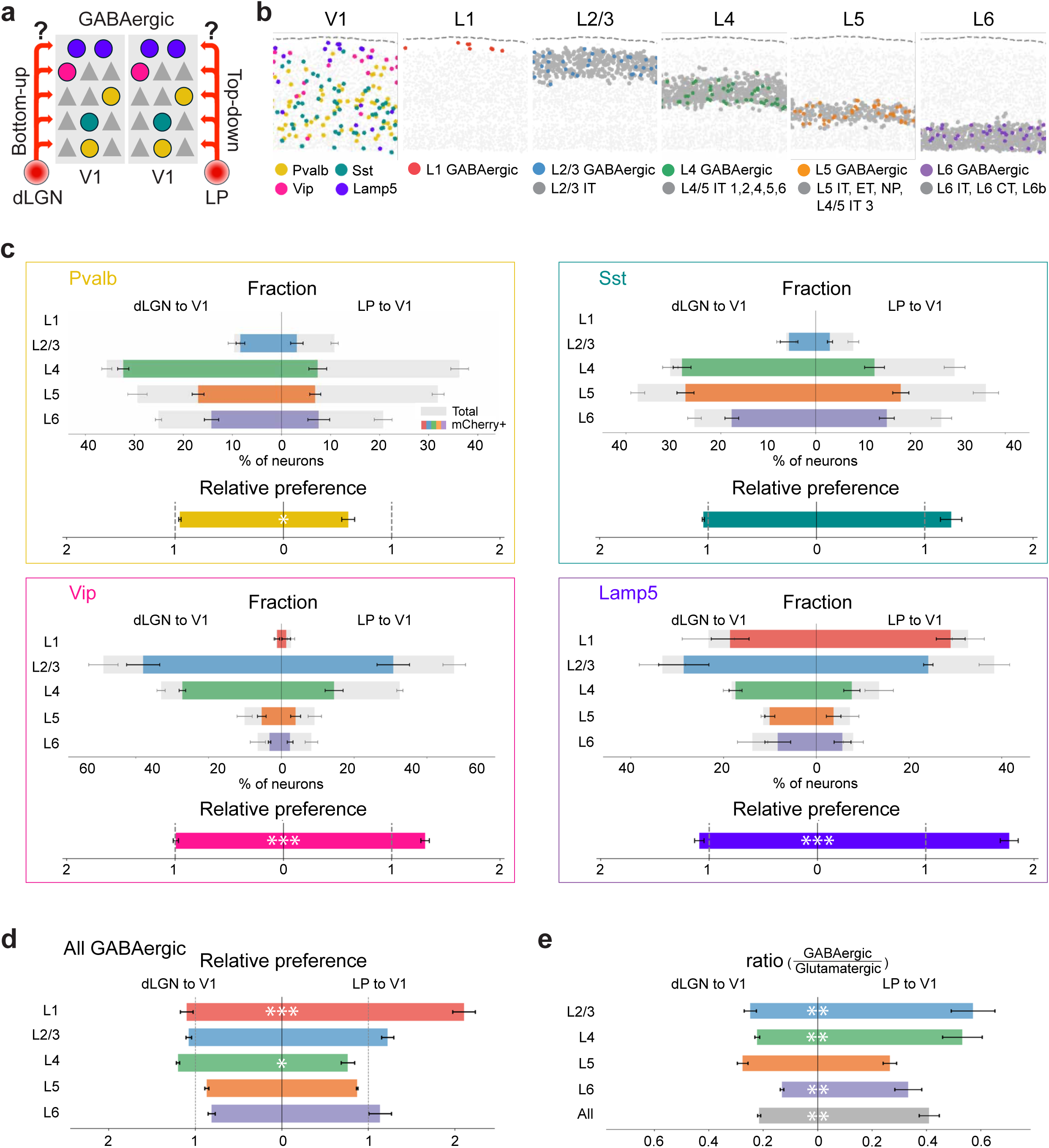
Distinct patterns of GABAergic V1 neurons targeted by dLGN and LP. **a**, Schematic illustrating the question: which molecularly defined GABAergic neurons are targeted by bottom-up dLGN input or top-down LP input? **b**, Spatial organization of GABAergic neurons in an example V1 section. Left, spatial distribution of GABAergic subclasses; colored dots indicate neurons belonging to the subclasses listed below, and all other neurons are shown in light gray. Right, spatial distribution of GABAergic neurons assigned to cortical layers (left to right, L1, L2/3, L4, L5 and L6). Colored dots indicate GABAergic neurons assigned to each layer, and dark gray dots indicate glutamatergic anchor cells used for proximity-based layer assignments. The subclass and supertype identities of the glutamatergic anchor cells used for each layer are indicated below. **c**, Top, fractions of mCherry^+^ neurons across cortical layers for individual GABAergic subclasses in V1 following dLGN or LP injections. Bottom, relative preference values for the corresponding subclass. Gray bars indicate the fraction of neurons in each layer out of all neurons of that subclass. Colored bars indicate the fraction of mCherry^+^ neurons in each layer out of all neurons of that subclass. Relative preference was calculated as the mCherry⁺ fraction of neurons in each subclass normalized to the mCherry⁺ fraction of all GABAergic neurons. Significance is shown only for relative preference. Significance was determined using two-sided Welch’s t-tests on log-transformed data, adjusted for multiple comparisons using FDR correction. **d**, Relative preference of GABAergic neurons (all subclasses combined) across cortical layers for dLGN and LP inputs to V1. Relative preference was calculated as the mCherry⁺ fraction of GABAergic neurons (across all subclasses) in each layer, normalized to the mCherry⁺ fraction of all GABAergic neurons. Significance was determined using two-sided Welch’s t-tests on log-transformed data, adjusted for multiple comparisons using FDR correction. **e**, GABAergic-to-glutamatergic ratios of neurons targeted by dLGN and LP inputs, shown for each layer (colored) and across all layers (gray; see Methods). Significance was determined using two-sided Welch’s t-tests on log-transformed data, adjusted for multiple comparisons using FDR correction, except the overall ratio, which was not adjusted for multiple comparisons. dLGN to V1: n = 4, LP to V1: n = 5. Bars represent mean ± s.e.m. across animals. \**P* < 0.05, \*\**P* < 0.001, \*\*\**P* < 0.0001. Full details on statistical models and *P* values are provided in Supplementary Table 4.

Similar to glutamatergic neuron innervation pattern above, the dLGN targeted a higher fraction of V1 GABAergic neurons than LP (dLGN: 75.55 ± 1.91%; LP: 40.42 ± 4.60%; *P* = 0.0006), across three of the five major subclasses, namely Pvalb, Sst and Vip (Extended Data Fig. 8a). However, the two projections showed distinct relative preferences for GABAergic subclasses and their laminar location. LP showed a stronger relative preference for Vip and Lamp5 neurons compared to the dLGN, two subclasses which are enriched in the superficial layers of V1 (Fig. 2c). The dLGN showed a stronger relative preference for Pvalb neurons and specifically for those located in layer 2/3 and layer 4 (Fig. 2c; Extended Data Fig. 8c). Consistent with these subclass and layer-specific differences, dLGN and LP projections exhibited distinct laminar preference among their targeted GABAergic neurons (Extended Data Fig. 8b and Fig. 2d; *F_4,35_* = 18.83, *P* = 2.44 × 10^-8^ for relative preference). No significant differences in supertype preference within subclasses were detected between dLGN and LP targets (Extended Data Fig. 7).

We confirmed these results with the Cre-On approach in which we conditionally transfected dLGN core neurons using the Crh-Cre mouse line and LP neurons using the Calb2-Cre mouse line. The overall composition of V1 glutamatergic and GABAergic subclasses and supertypes targeted by Crh-Cre and Calb2-Cre neurons was similar to the innervation pattern using the Cre-Off approach described above (Extended Data Fig. 9, 10). Consistent with past studies indicating that the dLGN core neurons show a reduced innervation of the superficial layers compared to dLGN shell neurons^30^, we observed a tendency for a reduced innervation of L2/3 IT neurons by Crh-Cre neurons (Extended Data Fig. 9).

The TransA-MERFISH datasets revealed that the dLGN and LP each target a distinct pattern of molecularly defined V1 glutamatergic and GABAergic neurons with distinct laminar distributions. To determine whether these two inputs also differ in the proportion of targeted glutamatergic and GABAergic neurons, we calculated the ratio between the mCherry⁺ GABAergic and glutamatergic neurons for each layer, excluding L1, which lacks glutamatergic neurons. Overall GABAergic-to-glutamatergic ratio was almost twice as large for LP compared to dLGN targets (dLGN: 0.22 ± 0.01; LP: 0.41 ± 0.04; *P* = 0.003, two-sided Welch’s t-test). Specifically, LP targets exhibited higher GABAergic-to-glutamatergic ratios in L2/3, L4 and L6, with no difference observed in L5 (Fig. 2e). These data demonstrate a strong bias toward GABAergic neurons in the top-down LP input compared with the bottom-up dLGN input.

Taken together, these experiments reveal the distinctive innervation patterns of V1 by the dLGN and LP and highlight three particularly salient attributes of the innervation by LP: A strong relative preference for the L5 ET glutamatergic subclass, while disfavoring supertype 0093 within that subclass; an innervation pattern of L4/5 IT supertypes that favors 0023 and 0025 and disfavors 0028, and an almost double GABAergic-to-glutamatergic ratio, compared to the dLGN. Are these attributes maintained in the innervation of HVCs by LP?

### LP input to HVCs reveals the visual hierarchy

V1 is surrounded by a system of HVCs located medially (MVC), laterally (LVC), and rostrally (AVC) relative to V1 (see methods for delineation; Extended Data Fig. 11). These HVCs are composed of the same major glutamatergic subclasses as V1, albeit with different relative proportions and each of these HVCs receives strong input from LP (Fig. 3a-d; Extended Fig. 12). How does the pattern of HVC neurons contacted by LP compare to that of V1 neurons contacted by LP? We addressed this question for each of the three regions of HVCs separately.

**Fig. 3.**
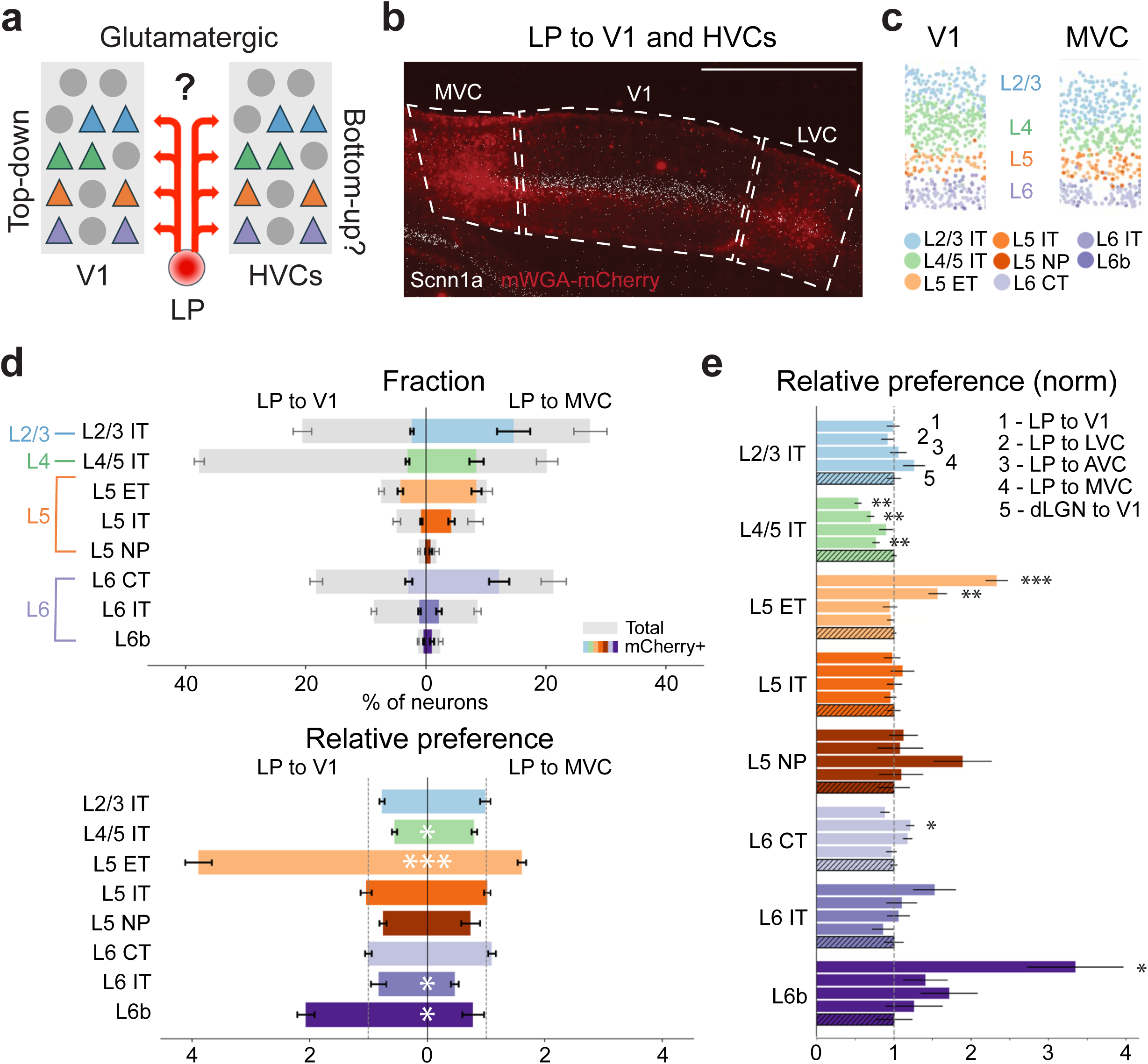
Patterns of glutamatergic neurons targeted by LP inputs to HVCs. **a**, Schematic illustrating the question: do LP inputs to HVCs target different patterns of molecularly defined glutamatergic neurons in HVCs as compared to in V1? **b,** Representative image showing anterogradely transferred mWGA-mCherry signal across visual cortical areas two weeks after LP injection with Scnn1a transcript distribution (white dots). MVC: medial higher visual cortex, LVC: lateral higher visual cortex. The higher expression of the Scnn1a transcript in V1 and the stronger mWGA-mCherry labeling in HVCs are used to delineate the boundaries between V1 and HVCs (see Methods). Scale bar, 1 mm. **c,** Example sections showing the spatial distribution of major glutamatergic subclasses in V1 and MVC. **d,** Top, fractions of mCherry^+^ neurons across glutamatergic subclasses in V1 (n = 5) and MVC (n = 5) following LP injections. Bottom, relative preference for each glutamatergic subclass in V1 and MVC targeted by LP inputs. Significance is shown only for relative preference. Significance was determined using two-sided Welch’s t-tests on log-transformed data, adjusted for multiple comparisons using FDR correction. Note that the relative preference of glutamatergic subclasses targeted by LP in MVC resembles the relative preference of glutamatergic subclasses targeted by the dLGN in V1 (Fig. 1g). **e,** Relative preference for glutamatergic subclasses targeted by LP inputs to V1 (n = 5), LVC (n = 5), AVC (n = 3) and MVC (n = 5), and dLGN input to V1 (n = 4). LP inputs to V1, LVC, AVC and MVC were measured in the same animals, with two animals lacking AVC data. Relative preference values for each subclass were normalized to those of dLGN input to V1. Asterisks denote significant differences in relative preference between LP inputs and dLGN inputs within each glutamatergic subclass. Significance was determined using two-sided Welch’s t-tests on log-transformed data, with FDR correction for multiple comparisons. Bars represent mean ± s.e.m. across animals. \**P* < 0.05, \*\**P* < 0.001, \*\*\**P* < 0.0001. Full details on statistical models and *P* values are provided in Supplementary Table 4. Note that the LP innervation patterns that differ the most from the dLGN to V1 innervation pattern are LP to V1 and LP to LVC.

The LP to MVC input targeted a pattern of glutamatergic and GABAergic neuronal subclasses that was distinct from the LP to V1 input pattern and that showed striking similarities to the dLGN to V1 input pattern. First, the LP input to MVC targeted an overall larger fraction of glutamatergic neurons than the LP to V1 input (across all glutamatergic subclasses except L5 NP and L6b; Fig. 3d). Second, it showed distinct relative preferences (*F_7,64_* = 7.69, *P* = 1.02 × 10^-6^) compared to the LP to V1 input, with a significantly increased preference for L4/5 IT and reduced preferences for L5 ET, L6b and L6 IT (Fig. 3d). Third, within the L4/5 IT supertypes, the LP input to MVC showed reduced preferences for supertypes 0023 and 0025 and an increased preference for supertype 0028; within the L5 ET subclass, it also showed an increased preference for supertype 0093 (Extended Data Fig. 13). As a consequence, the glutamatergic subclasses targeted by the LP to MVC input was very similar to the pattern targeted by the dLGN input to V1 (*F_7,56_* = 0.61, *P* = 0.75) (Fig. 3e). Fourth, the LP input to MVC showed distinct GABAergic targeting preferences compared to the LP input to V1 (*F_4,40_* = 3.25, *P* = 0.02): The relative preference for Lamp5 and Vip, and the underrepresentation of Pvalb, characteristic of the LP input to V1, were absent (Extended Data Fig. 12). Thus, the pattern of GABAergic neurons in MVC targeted by LP resembles that of V1 targeted by the dLGN (*F_4,35_* = 2.11, *P* = 0.10) (Extended Data Fig. 12). Finally, the GABAergic-to-glutamatergic ratio of LP-targeted MVC neurons (0.20 ± 0.01) was lower than that of LP-targeted V1 neurons (*P* = 0.0004) and similar to that of dLGN-targeted V1 neurons (*P* = 0.90) (Fig. 4a). Together, this analysis reveals that the LP to MVC input is strikingly distinct from the LP to V1 input and, instead, carries a clear bottom-up, dLGN-to-V1-like “signature”.

**Fig. 4.**
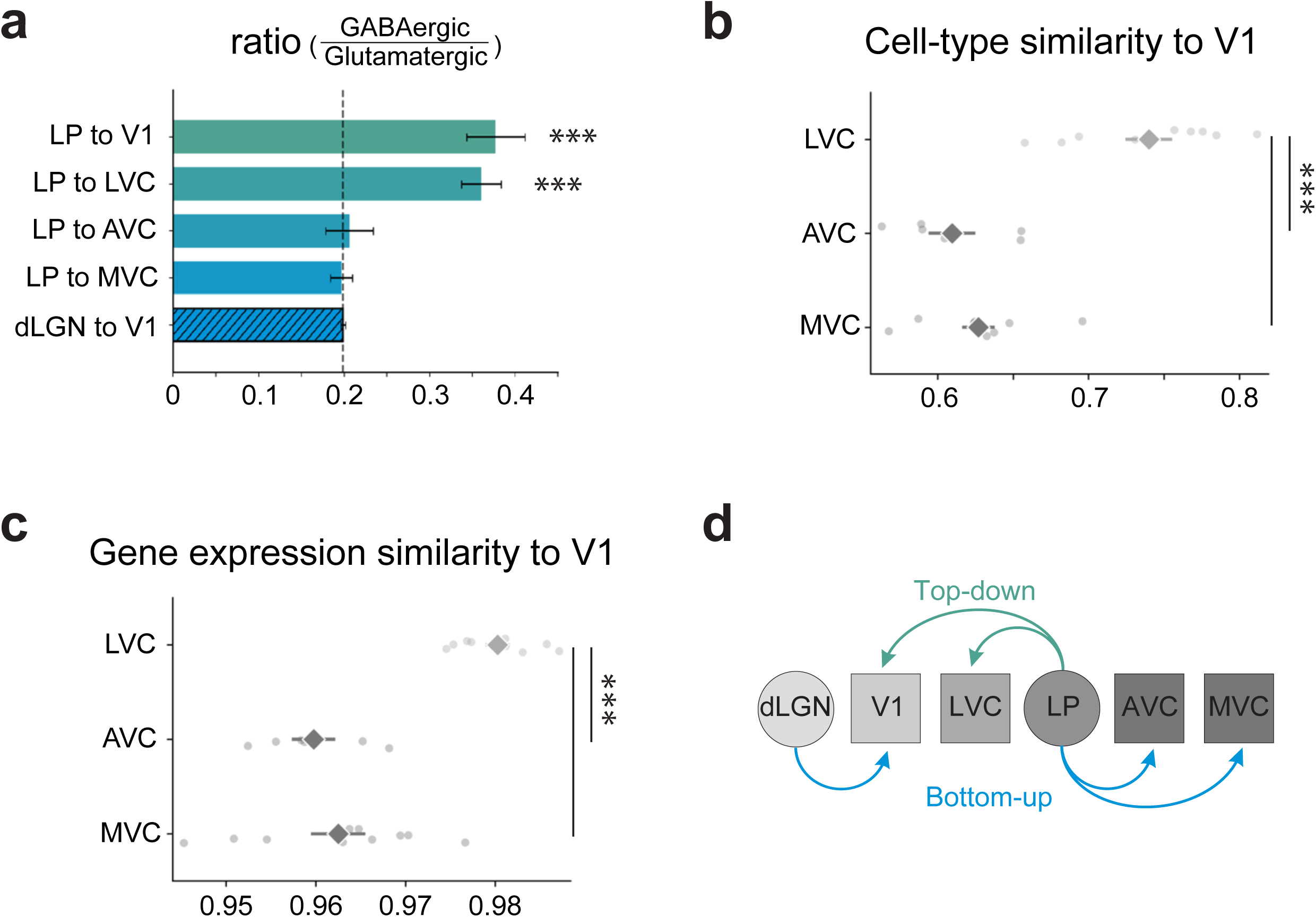
Hierarchy among HVCs. **a**, GABAergic-to-glutamatergic ratios of neurons targeted by LP inputs to V1 (n = 5), LVC (n = 5), AVC (n = 3) and MVC (n = 5), and dLGN input to V1 (n = 4). Asterisks denote significant differences between LP inputs to either visual area and dLGN input to V1. Significance was assessed using two-sided Welch’s t-tests on log-transformed data, with FDR correction for multiple comparisons. Note that LP to V1 and LP to LVC are both significantly different from dLGN to V1. **b**, Supertype composition similarity between V1, and LVC (n = 10), AVC (n = 6) and MVC (n = 10). Similarity was quantified using the Bray–Curtis metric (see Methods). LVC, AVC and MVC were measured in the same animals, with four animals lacking AVC data. Significance was determined using standard two-sided t-tests on logit-transformed data, with FDR correction for multiple comparisons. **c**, Bulk gene expression similarity between V1 and LVC (n = 10), AVC (n = 6) and MVC (n = 10). Similarity was quantified using Spearman’s rank correlation coefficient (see Methods). Significance was determined using standard two-sided t-tests on Fisher’s Z-transformed data, with FDR correction for multiple comparisons. **d**, Schematic illustrating the hierarchy among thalamic nuclei and visual cortical areas based on mapping the innervation pattern of thalamic nuclei onto the molecular taxonomy of cortical neurons. Bars represent mean ± s.e.m. across animals. \**P* < 0.05, \*\**P* < 0.001, \*\*\**P* < 0.0001. Full details on statistical models and *P* values are provided in Supplementary Table 4.

Next, we analyzed the pattern of AVC neurons targeted by LP. Like the LP input to MVC, the LP input to AVC had a dLGN-to-V1-like bottom-up signature. Accordingly, compared to the LP to V1 input, the LP to AVC input targeted a higher fraction of glutamatergic subclasses except L5 NP and L6b (Extended Fig. 9a). Furthermore, like the LP to MVC input, it showed a significantly increased preference for L4/5 IT, and within the L4/5 IT subclass it showed reduced preference for supertypes 0023 and 0025 and an increased preference for supertype 0028. It also showed a reduced preference for L5 ET yet, within that subclass, an increased preference for supertype 0093 (Extended Data Fig. 10a; Extended Fig. Data 10; Fig. 3e). Finally, the LP to AVC input also had a relatively low GABAergic-to-glutamatergic ratio (0.21 ± 0.03), like both the LP to MVC and the dLGN to V1 ratio (Fig. 4a). Thus, also the LP to AVC input has a dLGN-to-V1-like bottom-up signature.

Finally, we analyzed the innervation pattern of LVC by LP. In striking contrast to the innervation of MVC and AVC by LP, the innervation of LVC showed a pattern that was much closer to the top-down, LP to V1 input. Although LP targeted a higher fraction of glutamatergic subclasses in LVC (29.623 ± 2.828) compared to V1 (14.972 ± 1.676), it was significantly less than the fraction targeted in MVC (51.811 ± 2.883) and AVC (47.946 ± 1.642) (Extended Data Fig. 10b). Furthermore, similar to the LP input to V1, the LP input to LVC had a strong relative preference for L5 ET, yet disfavoring supertype 0093 within that subclass and, within the L4/5 IT subclass, showing increased preferences for supertypes 0023 and 0025 and reduced preference for supertype 0028 (Extended Data Fig. 10b; Extended Data Fig. 11; Fig. 3e). Notably, the GABAergic-to-glutamatergic ratio of the LP input to LVC was relatively large (0.36 ± 0.02), like the LP input to V1 (Fig. 4a). Thus, the LP to LVC input pattern shows an LP-to-V1-like, top-down signature.

Taken together, these data reveal two clearly distinct patterns in the innervation of HVCs by LP: a “dLGN-to-V1-like”, bottom-up pattern in MVC and AVC and an “LP-to-V1-like” top-down pattern in LVC. This pattern was corroborated with the Cre-On approach using the Calb2-Cre mouse line (Extended Data Fig. 13).

To determine whether these two distinct innervation patterns reflect similarities between visual cortical areas receiving either a bottom-up or a top-down input from LP, we compared the cell-type composition and bulk gene expression profiles derived from our MERFISH datasets across the three HVC subregions with those of V1. Interestingly, LVC showed greater similarity to V1 than did AVC or MVC at both the subclass and supertype levels (Fig. 4b; Extended Data Fig. 14a), with differences in relative cell-type proportions primarily driven by the glutamatergic populations (Extended Data Fig. 14a; these compositional differences are not related to the observed differences in relative targeting preferences, which are normalized to the underlying cell-type proportions). Likewise, bulk gene expression profiles were more similar between LVC and V1 than between V1 and either AVC or MVC (Fig. 4c; Extended Data Fig. 14b). Thus, the distinct innervation patterns by LP predict cellular and transcriptional similarities across HVCs.

The bottom-up and top-down input structure laid out here allows us to outline a hierarchy among thalamic nuclei and HVCs in which, with the dLGN assigned to the bottom of the hierarchy, V1 and LVC are located between the dLGN and LP, and AVC and MVC are on top of the hierarchy (Fig. 4d). Remarkably, this hierarchy, based on mapping the innervation pattern of thalamic nuclei onto the molecular taxonomy of cortical neurons, closely matches the hierarchy of visual areas based on independent criteria ^12,13^.

## Discussion

Single-cell transcriptomics has uncovered the cellular makeup of the mammalian visual cortex with unprecedented granularity. While compiling this parts list represents a major advance in our understanding of cortical structure, how these molecularly diverse cell types are integrated into the canonical cortical circuit subserving vision remains only rudimentarily understood. To bridge this gap, we integrated anterograde transsynaptic tracing with spatial transcriptomics (TransA–MERFISH), an approach that enables high-throughput identification of cortical neurons receiving direct thalamic input together with detailed cell-type classification, a combination not previously achievable. Specifically, we established a set of new virogenetic reagents, as well as streamlined histology and imaging protocols to apply TransA-MERFISH to thalamocortical studies. Using this high-throughput approach, we identified two basic thalamic innervation patterns of cortical neurons across primary, medial, anterior, and lateral visual areas: a dLGN-to-V1–like pattern and an LP-to-V1–like pattern, referred to as bottom-up and top-down, respectively. These two innervation patterns, or signatures, differ not only in the molecular subclasses and supertypes of the targeted cortical neurons but also in the ratio of innervated GABAergic to glutamatergic neurons. These signatures reveal a hierarchy of visual areas.

These findings highlight the critical importance of fine-scale molecular categories of neurons, such as supertypes, in understanding differences between thalamic innervation patterns. For example, a main difference in the innervation of V1 by the dLGN and LP is their distinct relative preferences for glutamatergic supertypes in L4/5 IT (supertypes 0023, 0025 and 0028) and L5 ET (supertype 0093). It is possible that additional data with greater statistical power will reveal differences in thalamic innervation patterns even among finer molecular categories, like clusters. Irrespective, the demonstration that supertypes are differentially integrated into the canonical cortical circuit lends support to the idea that molecularly defined supertypes within a given subclass are functionally distinct^31^.

Our data also demonstrate that molecularly distinct neuronal subsets within a given thalamic nucleus follow the overall cortical innervation pattern of that nucleus, but for a few specific differences in targeting preferences. The dLGN core-specific Crh-Cre neurons, for example, show reduced preference for V1’s L2/3 and L5 IT neurons compared to the general dLGN populations. Similarly, Calb2-cre neurons in LP show enhanced preference for L6b across all visual areas, compared to the general LP population. Thus, a complete understanding of thalamic innervation patterns to the cortex not only necessitates a granular definition of molecular types in the cortex but may also benefit from a commensurate categorization of thalamic neuron types^32,33^.

The shared cell-type targeting signatures of bottom-up and top-down thalamic inputs, irrespective of their thalamic nucleus of origin or their target cortical area, suggest common principles underlying the organization of these two information streams across cortical areas. While the present study focuses on thalamocortical inputs, future experiments comparing bottom-up and top-down signatures across thalamocortical and corticocortical pathways will help determine whether these principles are specific to thalamocortical inputs or extend more broadly across cortical circuits.

These findings also show that the pattern of cortical neurons targeted by thalamic top-down projections exhibit larger GABAergic to glutamatergic ratios than bottom-up projections. This difference does not automatically imply an increased inhibition to excitation ratio in cortex upon activation of top-down projections. In fact, top-down projections show a larger relative preference for the VIP GABAergic subclass compared to bottom-up projections, a type of inhibitory neurons that, by targeting other GABAergic neurons, contributes to disinhibition^34,35^. In addition, bottom-up projections show a larger relative preference for Pvalb GABAergic neurons, an inhibitory subclass providing strong somatic inhibition to pyramidal cells^36,37^. Consistent with the idea of a possible shared bottom-up and top-down signatures across thalamocortical and corticocortical pathways, electrophysiological and anatomical studies have shown a stronger excitation and innervation of Pvalb GABAergic neurons by bottom-up projections from V1 to HVC compared to top-down projections from HVC to V1^38,39^.

The mWGA-mCherry anterograde transynaptic transfer reveals the presence of connections, not the strength of the synapse. On one hand, the enhanced relative preference for certain glutamatergic and GABAergic neuron subclasses by the dLGN and LP observed here parallels findings using electrophysiological approaches^18,40–42^. On the other hand, we also show anterograde transfer between the thalamus and cortical targets in which synaptic physiology indicates weak transmission, for example between LP and layer 4 glutamatergic neurons in V1^20,43^. The presence of these connections, however, have been clearly reported by studies using anatomical techniques, like retrograde transsynaptic rabies-based protocols^44^, thus further validating our approach.

The bottom-up and top-down signatures of molecularly defined cortical neurons targeted by the thalamus allow us to define a hierarchy of visual areas in which V1 and LVC are positioned between the dLGN and LP, whereas AVC and MVC occupy the top of the hierarchy. Interestingly, this hierarchy closely matches previously established hierarchical orders based on different criteria^12,13,45^. Furthermore, while recent work has shown that gene expression and cell-type composition vary across cortical areas^9,10,46–51^, we find that similarities in these features across visual cortical areas align with their relative positions within the hierarchy. The convergence of these independent approaches demonstrates how molecular, cellular, and circuit-level features are orchestrated to establish cortical hierarchy.

## Methods

### Animals

Experiments were conducted in accordance with the regulations of the Institutional Animal Care and Use Committee of the University of California, San Francisco. Mice were maintained on a 12-h light/12-h dark cycle with regular housing conditions and standard access to drink and food. Mice 8-10 weeks of age were used for stereotaxic injections followed by MERFISH experiments 2-3 weeks post injections. No statistical methods were used to predetermine sample size. Mice of either sex and of the following genotypes were used (Cre driver lines were maintained as homozygous in the parental strains): Emx1-Cre (JAX #005628), Calb2-IRES2-Cre-D (JAX #028532), Crh-Cre (GENSAT; MMRRC #036507-UCD; a generous gift from Dr. Jamie Maguire, Tufts University); Sst-IRES-Flp (C57BL/6J) (JAX #031629). No statistical methods were used to predetermine sample size. The mice were randomly chosen. The experimenter was not blind to the experimental conditions.

### Adeno-associated viruses

The following adeno-associated viruses (AAV) were used:

AAV1.hSyn.EGFP (Addgene #50465, titer of 1.9 x 10^13^ genome copies/mL)

AAV9.EF1α.FAS.mWGA.mCherry (titer of 1.0 x 10^13^ genome copies/mL)

AAV9.CAG.Flex.mWGA.mCherry (titer of 1.0 x 10^13^ genome copies/mL)

AAV9.EF1a.DIO.hChR2(H134R)-eYFP.WPRE.hGH (Addgene #20298 titer of 2.6 x 10^13^ genome copies/mL)

AAV8.EF1a.Coff/Fon-BFP (Addgene #137131, titer of 1.6 x 10^13^ genome copies/mL)

### Stereotaxic injections

Mice were anesthetized with 1-2% isoflurane and received subcutaneous buprenorphine and carprofen for analgesia. Their body temperature was maintained at 37 °C using a feedback-regulated heating pad (FHC, 40-90-8D). A small craniotomy was performed. Viral suspensions were loaded into a beveled glass pipette and injected using either a hydraulic manipulator (Narashige, MO-10) or a micropump (WPI, UMP-3). To minimize viral backflow during pipette withdrawal, the pipette was left in place for at least 10 min following thalamic injections and 5 min following cortical injections. After virus injection, the skin was sutured with 6-0 nylon suture (Ethilon 1698G). The coordinates of the injection sites and the volumes of the injected viral suspension are detailed below:

dLGN: 2.4 mm lateral to the midline, 2.3 mm posterior to the bregma, 2.6 mm below the pia

V1 (for GFP expression): 2.6 mm lateral to the midline, 3.5 mm posterior to the bregma, 0.3mm below the pia

V1 (for ChR2 expression): 2.6 mm lateral to the midline, 3.5 mm and 3.0 mm posterior to the bregma, 0.3 and 0.6 mm below the pia

To express mWGA-mCherry in Emx1-Cre mice, ∼10 nl of AAV9.EF1ɑ.FAS.mWGA.mCherry (diluted to titer of 2.5 x 10^12^ genome copies/mL) was injected. To express mWGA-mCherry in Calb2-cre and Crh-cre mice, ∼20 nl of AAV9.CAG.Flex.mWGA.mCherry (titer of 1.0 x 10^13^ genome copies/mL) was injected. To express GFP in V1 of Emx1-cre mice, ∼10 nl of AAV1.hSyn.EGFP (titer of 1.9 x 10^13^ genome copies/mL) was injected 6 weeks after dLGN injection of AAV9.EF1α.FAS.mWGA.mCherry and 2 weeks before tissue extraction.

To verify the monosynaptic transfer of mWGA from dLGN to V1, in Crh-Cre mice, ∼150 nl of AAV9.CAG.Flex.mWGA.mCherry (titer of 1.0 x 10^13^ genome copies/mL) and ∼150 nl AAV9.EF1a.DIO.hChR2(H134R)-eYFP.WPRE.hGH (titer of 2.6 x 10^13^ genome copies/mL) were co-injected in dLGN 3-4 weeks before being perfused for slice electrophysiology. In Crh-cre; Sst-Flp mice, 2 weeks after injection of ∼150 nl of AAV9.CAG.Flex.mWGA.mCherry and ∼150 nl AAV9.EF1a.DIO.hChR2(H134R)-eYFP.WPRE.hGH into dLGN, AAV8.EF1a.Coff/Fon-BFP (titer of 1.6 x 10^13^ genome copies/mL) was injected at V1 at 2 sites with 2 depths, and ∼150 nl virus was injected at each depth. After 10-14 days, mice were perfused for slice electrophysiology.

### Slice recording

Mice (P50–P75) injected with viruses as described above (see “Stereotaxic injections”) were anesthetized by intraperitoneal injection of ketamine and xylazine (100 mg/kg and 10 mg/kg, respectively) and transcardially perfused with ∼5 ml cold (0–4 °C) modified artificial cerebrospinal fluid (ACSF) perfusion solution (in mM: 225 sucrose, 119 NaCl, 2.5 KCl, 1 NaH_2_PO_4_, 26.2 NaHCO_3_, 1.25 D-glucose, 3 kynurenic acid, 1 Na-ascorbate, 4.9 MgCl_2_ and 0.1 CaCl_2_, saturated with 95% O_2_/5% CO_2_). Following rapid decapitation, whole brains were dissected and transferred into ice-cold oxygenated ACSF cutting solution (in mM: 110 choline chloride, 2.5 KCl, 0.5 CaCl_2_, 7 MgCl_2_, 1.3 NaH_2_PO_4_, 1.3 Na-ascorbate, 0.6 Na-pyruvate, 20 glucose, and 26.2 NaHCO_3_, saturated with 95% O_2_/5% CO_2_, pH 7.4, 310-315 mOsm). Coronal slices of 250–300 μm were prepared in a cold cutting solution with a Leica VT1200S vibratome. Slices containing V1 were incubated in a submerged chamber at 34 °C for 30 min within ACSF recording solution (in mM: 125 NaCl, 2.5 KCl, 2 CaCl_2_, 1 MgCl_2_, 1.3 NaH_2_PO_4_, 1.3 Na-ascorbate, 0.6 Na-pyruvate, 20 glucose, and 25 NaHCO_3_, saturated with 95% O_2_/5% CO_2_, pH 7.4, ∼310 mOsm), and subsequently maintained at room temperature (∼22 °C) until recording. Whole-cell recordings were performed in ACSF recording solution (saturated with 95% O_2_/5% CO_2,_ perfused at 3ml/min) at 31–32 °C. For whole-cell recordings, we used either a K^+^-based pipette solution containing in mM: 143 K-gluconate, 10 HEPES, 1 EGTA, 2.5 MgCl_2_, 4 Mg-ATP, 0.3 Na-GTP, 10 Na_2_-phosphocreatine (295 mOsmol, pH 7.35) or a Cs^+^-based pipette solution containing in mM: 115 Cs-methanesulfonate, 10 HEPES, 1 EGTA, 1.5 MgCl_2_, 4 Mg-ATP, 0.3 Na-GTP, 10 Na_2_-phosphocreatine, 2 QX-314-Cl, 10 BAPTA-tetracesium (295 mOsmol, pH 7.35). Membrane potential was not corrected for the liquid junction potential. V1 neurons were visualized using DIC infrared video-microscopy with a water immersion objective (40×, 0.8 NA) on an upright microscope (Olympus BX51WI) with an IR CCD camera (Till Photonics VX44). Slices were examined under a 4× objective before recording to determine whether ChR2-EYFP and mWGA-mCherry were expressed in the dLGN. In slices with robust expression, whole-cell voltage-clamp recordings were selectively performed on V1 pyramidal neurons with mCherry fluorescence and neighboring mCherry-negative neurons, and V1 inhibitory Sst neurons labeled with BFP in Crh-Cre; Sst-Flp.

AMPA receptor-mediated EPSCs were recorded at the IPSC reversal potential (∼−65 mV). To verify thalamocortical monosynaptic connectivity, isolated monosynaptic AMPA receptor-mediated EPSCs using a modified sCRACM approach in the presence of tetrodotoxin (TTX; 1 μM; Tocris 1069), 4-aminopyridine (4-AP; 1.5 mM; Abcam ab120122), and tetraethylammonium (TEA; 1.5 mM; Abcam ab120275). EPSCs were low-pass filtered at 4 kHz and acquired at 10 kHz using a Multiclamp 700B or 700A amplifier and digitized at 10 kHz with a Digidata 1440A under the control of Clampex 10.2 (Molecular Devices). Data was analyzed offline using Clampfit 10.2 (Molecular Device) and Matlab. To photostimulate ChR2-expressing thalamocortical axons, blue light (470 nm) was delivered using a collimated LED and a T-Cube LED Driver (Thorlabs) through the fluorescence illuminator port and a 40× objective. Light pulses (5ms or 10 ms; 5.5 mW/mm^2^) were delivered with a 30 s inter-stimulus interval.

Monosynaptic EPSC amplitudes were measured from an average of 8-10 sweeps. For the individual sweep, the baseline was calculated as the median of the 10 ms region preceding LED driver ON. We considered a V1 cortical neuron to receive thalamic input when the average EPSC amplitude was at least three times the baseline standard deviation.

### MERFISH gene-panel selection and validation

To discriminate transcriptionally distinct cell types, we designed a panel of 300 genes following a computational pipeline established from previous MERFISH studies for the mouse brain^1,2^. Among the 300 genes, 181 are selected from established marker genes of visual cortical cell types identified in single-cell RNA sequencing (scRNA-seq) studies ^3–5^, with a focus on neuronal cell types. To further enhance discrimination between different GABAergic neuronal types, we selected the top 50 genes that provide the highest mutual information in defining GABAergic neuronal clustered in the Allen Institute Mouse V1 SMART-seq dataset^4^, using a previously reported approach^1^. Due to overlap between these 50 genes and the previous 180 genes, 27 new genes were identified. In addition, we identified a panel of 36 new genes based on pairwise differential expression analysis of the V1 SMART-seq dataset across all neuronal clusters ^1^. Briefly, for each neuronal cluster pair, in both directions, candidate genes are identified if they exhibit at least a twofold difference in mean expression, be expressed in at least 40% of cells in the foreground cluster, show greater than threefold enrichment in the fraction of expressing cells relative to the background cluster, and reach statistical significance (P < 0.05; ANOVA test on log-transformed expression values). These candidate genes were then selected via a greedy algorithm that iteratively added markers to ensure every cluster pair was distinguished by at least two genes. The panel also included the gene *Cre*. Finally, due to historical reasons, 55 additional genes originally designed to discriminate against dLGN neuronal cell types were included in the panel. All the genes in the panel were included in the analysis.

In addition to the 300-gene MERFISH panel, we also imaged two reporter genes, *mCherry* and *tdTomato*. Because the coding sequences of *tdTomato* and *mCherry* are highly similar, either probe could be used to identify *mCherry* expression in cells transduced with AAV mWGA-mCherry virus. Due to the potential high copy number of *mCherry* transcripts in transduced cells, both reporter genes were imaged in a separate, sequential round of FISH imaging following the MERFISH run.

Encoding probes for the 300-gene MERFISH panel and the sequentially imaged reporter genes were constructed by Vizgen Inc using a commercial pipeline.

While the MERFISH gene panel was designed based on earlier Allen Institute scRNA-seq studies focused on visual cortex, cell identities in this study were assigned using the updated whole-brain taxonomy^6^ (CCN20230722). To evaluate how well the panel resolves cell types defined by this updated taxonomy, we performed in silico validation using the corresponding scRNA-seq dataset. Expression matrices from isocortical regions were loaded and restricted to genes included in the MERFISH panel. Gene expression values were library-size normalized to 10,000 counts per cell and log-transformed. We then trained a Random Forest classifier using only MERFISH panel genes to predict cell-type identity at the level of class, subclass, and supertype. Models were trained on 80% of the data and evaluated on a held-out 20% test set using stratified sampling. The classifier was implemented with 100 trees and class weights balanced to account for unequal cell-type representation. Classification accuracy was computed, and performance was further assessed using row-normalized confusion matrices to quantify separability between cell types. Validation was performed for major cell classes, neuronal subclasses, and glutamatergic and GABAergic supertypes separately, using inclusion criteria consistent with downstream analyses (see “Cell type assignment and QC”).

### Tissue preparation for MERFISH

#### Tissue collection

All tools and work surfaces were treated with RNase decontamination solution (RNaseZap, Invitrogen) and 70% ethanol, or exposed to UV light, to minimize RNase contamination following standard RNA handling procedures. Brain tissue for MERFISH was collected two weeks after injection of AAV9.EF1α.FAS.mWGA.mCherry (Cre-Off) or three weeks post injection of AAV9.CAG.Flex.mWGA.mCherry (Cre-On) into the thalamus. To harvest tissue, mice were anesthetized with isoflurane. The brain was rapidly extracted and immersed in 4 mM Ribonucleoside Vanadyl Complex (RVC) (NEB, S1402S) in 1× PBS in a 60mm x 15mm Petri dish (CELLTREAT, 229663). The immersed brain was bisected along the midline, and the hemisphere that received virus injection was retained. The retained hemisphere was further trimmed by removing the tissue anterior to bregma and the ventral half to reduce the cross-sectional area, allowing multiple sections to be mounted on a single MERFISH slide. The trimmed tissue block was immersed in 4% paraformaldehyde (PFA) in 1× PBS for ∼3 h at room temperature, and then transferred to 30% sucrose in 4 mM RVC in 1× PBS solution at 4 °C. This fixation duration was sufficient to preserve mCherry localization, preventing diffusion after cryostat sectioning (which disrupts mCherry imaging quality), without affecting transcript detection, and avoiding variability associated with PFA perfusion due to potential vascular clogging. Once the tissue block sank to the bottom in the 30% sucrose solution overnight at 4 °C, it was transferred to a Petri dish with OCT (Epredia, 22-110-617) to remove residual sucrose solution. The tissue block was placed in OCT in a plastic mold (Sakura Finetek, 10690461) and immediately frozen on powdered dry ice. After freezing, the sample was placed in a sealed plastic bag and stored at -80 °C until cryostat sectioning.

#### Cryosectioning

Cryostat sectioning of tissue blocks was performed using either a Leica CM1950 or a CryoStar NX70 cryostat (Epredia). All tools and cryostat surfaces were treated with an RNase decontamination solution (RNaseZap, Invitrogen) and 70% ethanol, or exposed to UV light, prior to sectioning. Two sets of coronal sections were prepared. The first set was used for immunohistochemistry to assess mWGA–mCherry expression at the injection site prior to the MERFISH workflow. For this set, sections were cut at a thickness of 40 μm. Every other section, starting anterior to the thalamus before the mWGA-mCherry signal was detectable and continuing to the posterior cortex (except when MERFISH sections were collected in between), was mounted onto Superfrost Plus slides (Fisherbrand, 12-550-15) for subsequent staining (see Immunohistochemistry section below for details). The second set consisted of 12 μm sections for MERFISH, collected at ∼400 μm intervals across four anterior–posterior positions starting at ∼2.3 mm posterior to bregma. For each MERFISH slide (Vizgen), four sections were mounted, and multiple slides were collected per tissue block for independent RNA/protein validation and as backups. Across slides, sectioning was repeated at the same coordinates with a ∼12 μm offset (e.g., 2.3, 2.7, 3.1, and 3.5 mm posterior to bregma). In a subset of experiments, MERFISH slides were coated with Poly-D-Lysine (Gibco, A3890401) prior to cryosectioning to promote section adhesion.

During cryosectioning, quality control was performed by checking the mWGA-mCherry expression at the injection site using an upright microscope (Olympus, MVX10). Tissue blocks with weak or off-target expressions, such as LP injections leaking into dLGN or vice versa, were excluded from further experiments.

### MERFISH imaging with integrated antibody staining

Following cryosectioning, MERFISH slides with mounted sections were left in the cryostat for 1 h to promote section adhesion to the slide prior to further processing. Upon transferring to room temperature, sections were fixed with 4% PFA for 10 min, washed three times with 1× PBS, and then permeabilized in 70% ethanol at 4 °C overnight and stored in 70% ethanol at 4 °C for up to one month prior to further processing.

Samples that met quality criteria following immunohistochemistry (see the section below for details) were processed for MERFISH according to the manufacturer’s instructions for fresh and fixed frozen samples (MERSCOPE Fresh and Fixed Frozen Tissue Sample Preparation User Guide, 91600002). Briefly, samples underwent protein staining for mCherry, encoder hybridization, gel embedding, clearing, and staining for PolyT and DAPI prior to imaging. Protein staining for mCherry was performed using a rabbit anti-RFP primary antibody (Rockland, 600-401-379, 1:100) following the manufacturer’s protocol. In a subset of experiments, a modified protocol was used in which RNA anchoring was performed prior to gel embedding and clearing, followed by encoder hybridization, in accordance with the manufacturer’s instructions. This modification was implemented to enhance transcript retention and reduce experimental variability, while yielding results consistent with those obtained using the standard protocol.

The MERFISH imaging process was performed using the MERSCOPE instrument according to the manufacturer’s instructions (MERSCOPE Instrument User Guide, 91600001). Eight z stacks were captured over a thickness of 12 µm. After imaging, transcript decoding was performed using the Merscope proprietary software. The resulting transcript matrix containing the transcript identity and spatial coordinates, as well as images of DAPI, PolyT, and mCherry protein staining, were used for downstream analyses. MERFISH experiments with low RNA molecule detection (<15k RNA molecules per field of view; field of view defined as the single unit imaging field of 221.18 x 221.18 *µm*^2^) or poor mCherry staining quality were excluded.

### Immunohistochemistry

For each sample, 40 μm sections collected separately from MERFISH sections were immunostained for mCherry protein. Sections were fixed in 4% PFA for 15 min at room temperature, followed by three washes in 1× PBS (10 min each). Sections were then incubated in a blocking solution containing 5% goat serum and 0.5% Triton X-100 in PBS for 1 h at room temperature. After blocking, sections were incubated overnight at 4 °C in a solution containing 5% goat serum, 0.5% Triton X-100, and primary antibody (rabbit anti-RFP, Rockland, 600-401-379, 1:500) in 1×PBS. The following day, sections were washed three times in 1× PBS (10 min each) and incubated for 2 h at room temperature in a solution containing 5% goat serum, 0.5% Triton X-100, and secondary antibody (Alexa Fluor 568 goat anti-rabbit, Invitrogen, A11011, 1:500) in 1× PBS. Sections were then washed three times in 1× PBS (10 min each), air-dried at room temperature, and coverslipped with an antifade mounting medium containing DAPI (Vectashield, H-1500-10). After immunostaining, samples with weak or off-target mWGA-mCherry injections, such as LP injections leaking into dLGN or vice versa, were excluded from further experiments.

### Control experiment to exclude polysynaptic (secondary) transfer

In this control experiment, AAV9.EF1α.FAS.mWGA.mCherry (Cre-Off) was injected into the dLGN of Emx1-Cre mice. Six weeks after injections in the dLGN, AAV1.hSyn.EGFP was injected into the ipsilateral V1 (see “Virus injection” section for injection coordinates, volume and virus titer). Two weeks after V1 injection, mice were perfused with 5 mL 1×PBS followed by 15 mL 4% PFA. Brains were then extracted, fixed in 4% PFA overnight, cryoprotected in 30% sucrose overnight, and embedded in OCT for cryosectioning. 40 μm coronal sections spanning the dLGN and visual cortex in the anterior-posterior axis were collected. Sections were immunostained for mCherry and GFP protein following the same protocol (see “Immunohistochemistry” section), with the initial 4% PFA fixation step omitted. For GFP staining, a primary antibody against GFP (chicken anti-GFP, Aves Labs, NC9510598, 1:500) and a secondary antibody (Alexa Fluor 488 goat anti-chicken, Invitrogen, A-11039, 1:500) were used. Sections were imaged with a camera (Olympus, DP72) attached to an MVX10 stereoscope (Olympus). To test whether mWGA-mCherry labeling extends beyond one synapse downstream of thalamic neurons, we examined a circuit in which V1 neurons project to the retrosplenial cortex, a region that receives no direct thalamic input. Despite robust mWGA-mCherry labeling in V1, no labeled neurons were detected in retrosplenial cortex even up to three months after injection (Extended Data Fig. 3), indicating that mWGA-mCherry transfer across thalamocortical synapses remains monosynaptic over the time window examined.

### Validation of the Cre-Off strategy

As briefly described in the main text, to eliminate the retrograde update of AAV–mWGA-mCherry by corticothalamic neurons, we used two complementary genetic strategies to restrict mWGA-mCherry expression to thalamic neurons, including the Cre-Off configuration^7^, where AAV-Cre-Off-mWGA-mCherry was injected into the dLGN or LP of Emx1-Cre mice. Because Cre is expressed in all cortical glutamatergic neurons, including corticothalamic neurons, mWGA-mCherry expression was selectively suppressed in the corticothalamic neurons while remaining intact in thalamic neurons, thereby preventing retrograde cortical expression.

The specificity of this approach was validated using spatial transcriptomic imaging. Following AAV-Cre-Off-mWGA–mCherry injection into the dLGN, mWGA–mCherry protein was detected in thalamic neurons and in cortical neurons consistent with anterograde transsynaptic transfer, whereas mWGA–mCherry mRNA was restricted to thalamic neurons and undetectable in cortex (Extended Data Fig. 1), confirming the absence of retrograde cortical expression.

### Cell segmentation and transcript assignment

For each section, only the cortical area manually selected in MATLAB was used for cell segmentation and subsequent analyses. Regions with wrinkles or large bubbles that obscured imaging were excluded. Cells were segmented based on DAPI and PolyT staining using Cellpose^8,9^. Briefly, segmentation was performed on the maximum-intensity projection of multiple z-planes (3rd to 7th) using a manually curated pre-trained model. The resulting cell boundaries were propagated to z-planes above and below. After cell segmentation, transcripts were assigned to each cell based on their spatial coordinates, resulting in a cell-by-gene matrix. Cells with a mask area < 97.92 *µm*^3^ or > 3149.11 *µm*^3^ (three times the median volume of all cells)^1,6^, fewer than 10 genes detected, or fewer than 100 RNA molecules were excluded.

### Annotation of mWGA-mCherry-positive postsynaptic neurons in visual cortices

To determine which cells in the cortex received mWGA-mCherry transfer (or mCherry^+^), we performed the following analyses.

First, images from the mCherry channel were preprocessed in MATLAB to reduce noise and background. A 3 x 3 pixel median filter was applied to each of the 8 z-planes for denoising. After denoising, a mean-intensity projection image was generated. To correct for the uneven background in different cortical layers arising from the varying thalamocortical axon density and scattering, a rolling ball filter (ball radius = 50 pixels, corresponding to 5.4 *µm*) was used. The ball radius was designed to remove uneven background while retaining the punctated mCherry signal that was much smaller than the size of the soma.

Second, the average intensity of the mCherry signal for each cell within the region of interest was quantified and normalized to the local background. Because the mCherry signal was frequently enriched along the cell membrane (as illustrated in Fig. 1b), signal intensity was computed using a ring-shaped mask (15 pixels, corresponding to 1.62 µm in width) with the outer edge aligned to the boundary of the cell mask (inner ring). Local background intensity was estimated as the average mCherry signal within an external ring 25 pixels in width, with the inner edge positioned 10 pixels away from the cell mask boundary. The background-normalized mCherry intensity for each cell was defined as the mean intensity within the inner ring minus the mean intensity within the outer ring and was used for the subsequent analyses. The 15-pixel inner ring width was selected to preferentially capture signals near the membrane while minimizing inclusion of lower-intensity intracellular regions. Increasing the inner ring width led to increased variability in the mCherry⁺ population and reduced separation between mCherry⁺ and mCherry⁻ populations, as assessed by Gaussian mixture modeling.

Third, to identify mCherry^+^ cells, we performed a constrained two-component Gaussian mixture model (GMM). Specifically, for each animal, the intensity distribution of all cells in the manually selected cortical region of interest across multiple sections (up to 4 sections per animal) was fitted to two Gaussian components, corresponding to mCherry^-^ and mCherry^+^ populations. Two constraints were imposed to stabilize the fit and ensure biological interpretability. First, the mean of the mCherry- component was fixed at zero, consistent with background-subtracted intensity values. Second, the standard deviation of the mCherry^-^ component was constrained to lie within [0.5, 2.0] times the standard deviation computed from an estimated mCherry^-^distribution, which included the negative tail and the mirrored negative tail of the intensity distribution. Model parameters were learned using expectation–maximization until convergence. The fitted global model for each animal was applied to individual cells to compute posterior probabilities of mCherry^+^ positivity. The posterior probability of individual cells were then used to estimate the fraction of mCherry^+^ within each neuronal cell type.

### Cell type assignment and QC

To assign molecular identity to each cell, the cell-by-gene matrix was mapped onto the 10x Whole mouse brain taxonomy (CCN20230722)^6^ using hierarchical mapping in MapMyCells ( RRID:SCR_024672), a software package developed by the Allen Institute^10^. While all cells were assigned an identity at the class, subclass, supertype, and cluster levels, downstream analyses were focused on the class, subclass, and supertype levels. Several custom modifications were made to the MapMyCells pipeline: First, because our analyses were restricted to cortical regions, we mapped our data to a subset of the taxonomy containing only the neocortex and hippocampal formation. This was implemented by modifying the “nodes_to_drop” parameter in the mapping configuration. Second, we replaced the default marker-gene subsampling bootstrap with a count-thinning based robustness estimation (see below for details).

In the default MapMyCells workflow, each bootstrapping iteration randomly selects 50% of the marker genes to perform cell-type mapping. The final type assignment for each cell is determined by the dominant vote across 100 bootstrapping iterations, with the fraction of votes for the dominant type being the assignment robustness. This approach has two limitations. First, cell-type assignment can become unstable when highly informative genes are omitted, particularly for closely related cell types that rely on a small number of markers for discrimination. For example, if the distinction between two cell types relies primarily on a single gene, that gene has a 50% chance of being excluded in each bootstrap iteration, resulting in at least 50% chance of incorrect assignment and artificially reduced assignment robustness even when transcript detection is high quality. Second, the resulting robustness metrics reflect sensitivity of mapping to marker gene subsampling rather than the intrinsic quality of the cell’s transcriptomic profile and therefore is not ideal for cell-level quality control.

To avoid these issues, we first assigned each cell to a single cell type using the full marker gene set. We then quantified assignment robustness using a count-thinning bootstrap approach. Specifically, in each bootstrapping iteration, for each cell, the transcript counts of each gene were independently subsampled to 50% by randomly drawing from a binomial distribution with probability p = 0.5, thereby simulating technical variability in transcript detection while preserving the relative gene expression profile. Cell type assignment was then performed on the thinned count data. Assignment robustness was calculated as the fraction of votes for the dominant assigned type after 100 iterations. This robustness metric reflects the stability of cell type assignment under count perturbation and was computed at all levels of the cell type hierarchy in the taxonomy, including class, subclass and supertype. Cells with assignment robustness < 0.8 at the class level and < 0.8 at the subclass level were discarded from downstream analyses. We further removed a small fraction of neurons with misalignment between their assigned type and spatial location (see “Layer assignment” below for more details). These combined cell-level filtering criteria removed about 9% of all the neurons (67597 out of 73324 neurons remained in V1, 38834 out of 43096 neurons remained in HVCs).

In addition to the cell-level filtering, subclasses with very few assigned cells were excluded from downstream analyses. Specifically, cells from the same region (V1 or HVCs) were pooled within each experimental group (Emx1-Cre with dLGN injection, Emx1-Cre with LP injection, Crh-Cre and Calb2-Cre). Within each region, subclasses that consistently comprised >1% of the total cells within the corresponding class across groups were retained. This procedure yielded a common set of 8 glutamatergic and 5 GABAergic subclasses in both V1 and HVCs, collectively representing over 97% of all neurons that passed cell-level filtering.

Within each subclass, to ensure sufficient statistical power given low cell counts for some supertypes, we applied two criteria to determine which supertypes to include in the figures: First, a region-specific statistical floor was imposed to ensure sufficient sampling depth, with minimum cell count thresholds adjusted by region (V1: ≥50 GABAergic, ≥100 glutamatergic; HVCs: ≥35 GABAergic, ≥70 glutamatergic). Second, to account for the long-tailed distribution of rare cell types, we applied an adaptive cumulative filtering approach within each subclass, retaining the most abundant supertypes that collectively accounted for 98% of the subclass population, including the supertype that exceeded this threshold. These filtering procedures were performed on pooled datasets across all groups for each region, resulting in 29 glutamatergic supertypes in V1 and 34 in HVCs, and 30 GABAergic supertypes in V1 and 31 in HVCs. Neurons belonging to low-abundance supertypes that did not meet these criteria were nevertheless retained in the total mCherry⁺ population used for calculating overall fractions and relative preferences.

### Visualization of cell-type assignment using UMAP

To visualize the topological relationships between hierarchically assigned cell types, Uniform Manifold Approximation and Projection (UMAP) embeddings were generated. Cells that passed quality-control filters and were assigned to predefined glutamatergic and GABAergic subclasses were included. Raw transcript counts were normalized to a target sum of 10,000 counts per cell and log1p-transformed prior to dimensionality reduction. For the global embedding, all valid neurons across V1 and higher visual areas were combined. Principal component analysis (PCA) was performed on the normalized data, and a nearest-neighbor graph was constructed using the top 30 principal components. UMAP was then computed to visualize the overall organization of inhibitory subclasses and the continuous transcriptomic gradients across excitatory subclasses. To visualize supertypes within the L4/5 IT subclass, a separate UMAP embedding was generated for the L4/5 IT subclass. Cells were restricted to this subclass and to supertypes that passed filtering criteria. The count matrix for this subset was independently normalized and log-transformed, followed by PCA, nearest-neighbor graph construction, and UMAP embedding.

### Layer assignment

Each cortical neuron was assigned to a cortical layer based on its cell type assignment and spatial location. Since most of the glutamatergic subclasses in the cortex are layer-specific, the layer assignment for neurons belonging to these subclasses is straightforward. Specifically, in the visual cortex, L2/3 IT (L2/3 IT CTX) neurons were assigned to L2/3; L5 IT, L5 ET, and L5 NP neurons were assigned to L5; and L6 IT, L6 CT, and L6b were assigned to L6. L4/5 IT neurons occupy both L4 and L5. By examining the spatial overlap between L4/5 IT supertypes and L5 subclasses (L5 IT, ET, and NP), we observed a clear overlap between supertype L4/5 IT 3 and L5 but not the other L4/5 supertypes. Therefore, neurons of supertype L4/5 IT 1, 2, 4, 5, and 6 were assigned to L4, whereas neurons of supertype L4/5 IT 3 were assigned to L5. Note that L4/5 IT 4 neurons were observed only in HVCs and not in V1. To assign L4/5 IT neurons, we examined spatial overlap between individual L4/5 supertypes with all L2/3 subclasses and with L5 subclasses. As a result, supertypes L4/5 IT 2, 5 were assigned to L2/3, whereas supertypes L4/5 IT 1, 3, 4, 6 were assigned to L5.

After the layer assignment of these glutamatergic subclasses, two approaches were used to discard a small fraction of neurons showing clear disagreement between their spatial location and the assigned cortical layer. These neurons were also removed from further analyses. First, we removed neurons positioned at the inappropriate cortical depth relative to their neighboring neurons of the same layer. To do this, an estimated pia was manually traced for each cortical section using the Napari GUI in Python. A sliding window (108 *µm*) was applied along the pia’s length. For each cortical layer, the distance to the pia was calculated for all neurons within each sliding window. Neurons were identified as outliers if their distance-to-pia deviated substantially from neighboring neurons of the same layer, quantified using a modified z-score^11^, which is based on the median and median absolute deviation and is therefore robust to extreme values. It is defined as modified *Z* = |*x*_*i*_ − *M*|/(1.4826 × *median*|*x*_*i*_ − *M*|), where xᵢ is the distance-to-pia of an individual neuron, M is the median distance-to-pia of neurons assigned to the same layer within the window. The scaling factor 1.4826 rescales the median|x_i_ − M| to approximate the standard deviation under normality. The modified z-score is used to avoid extreme values from interfering with the mean and standard deviation. Neurons with a modified z-score > 3 were discarded. Second, isolated neurons that were far from neighboring neurons of the same layer were filtered out. Specifically, for each neuron, the number of same-layer neurons within a 65 *µm* radius was quantified. A neuron was discarded if there were < 3 same-layer neurons within this neighborhood.

The layer-assigned glutamatergic neurons were used as anchors for proximity-based layer assignment of the remaining neurons, including all GABAergic neurons. This customized approach does not rely on neurons’ absolute cortical depth for layer assignment, which can introduce noise due to variations in cortical thickness across different cortical areas.

First, a baseline anchor density was estimated for each cortical layer. Specifically, the number of anchor cells within a 65 µm radius was counted on a regular grid (54 µm spacing) spanning the cortical section. For each layer, maximal local density was estimated by averaging the densities of the top 5 most densely populated grid locations.

Second, for each neuron to be assigned, a neighborhood of 65 µm radius was defined. Within this neighborhood, the number of anchors that belong to each layer was quantified. Only layers with > 2 anchors in the neighborhood were considered. If a single layer met the criteria, the neuron was assigned to that layer. If multiple layers were eligible, the baseline anchor densities for these layers were compared, and the layer with the sparsest anchor cells was identified.

Anchor cells from denser layers were then randomly subsampled to match the density of the sparsest layer. Following this normalization, distances between the neuron and the remaining anchor cells were computed, and the neuron was assigned to the layer of its nearest anchor cell. Neurons initially assigned to L2/3 were subsequently evaluated for potential reassignment to L1, as described below.

Third, neurons initially assigned to L2/3 were re-evaluated for potential assignment to L1. Specifically, a convex hull around all L2/3 anchors was built. Neurons were reassigned to L1 if they were closer to the pia than the L2/3 hull boundary. This was determined by casting a ray from the pia towards the neuron; if the ray intersected with the L2/3 hull boundary after it encountered the neuron, then the neuron was reassigned to L1. Furthermore, if no layer met the assignment criteria, the neuron was assigned to L1, as neurons located at the boundaries of all anchor-defined layers are expected to have few neighboring anchor cells; however, this assignment was restricted to exclude cells likely belonging to deeper layers, including those with distances to the pia in the top 10% or those located within the L2/3 boundary but more than 108 *µm* below the hull, which were left unassigned.

### Cortical area identification and ROI selection

V1 borders were delineated in all mice. The borders of HVCs (AVC, MVC, and LVC) were delineated in mice with LP injections (including Emx1-Cre and Calb2-Cre mice). Several complementary approaches were used to delineate the borders of these cortical areas:

First, the morphology of the hippocampal formation was used to determine the anterior-posterior location relative to bregma for each section. Brain sections were manually registered to the Allen Adult Mouse Brain Atlas V3p1 using the ImageJ Aligning Big Brains and Atlases (ABBA) plugin. However, the resulting cortical area boundaries were often imprecise due to the limited spatial coverage of the sections (approximately one-third of the hemisphere), which hindered accurate estimation of section tilt along the mediolateral and dorsoventral axes.

Second, to refine the borders between V1 and HVCs for each animal, the *Scnn1a* gene expression pattern and the spatial distribution of mCherry^+^ cells were used. *Scnn1a* gene exhibited dense expression in L4 of primary sensory cortices and sparse expression in secondary sensory cortices^12,13^ (Fig. 3b), providing a clear molecular border between V1 and HVCs. Note that in our MERFISH data, *Scnn1a* expression was almost absent in MVC but detectable in LVC; however, *Scnn1a* transcript density in LVC was still substantially lower than V1 and was therefore readily distinguishable. In addition, in all experimental groups, the distribution mCherry^+^ cells was relatively homogeneous within V1 and distinct from HVCs, offering a complementary criterion to delineate the borders between V1 and HVCs (Fig. 3b). Note that due to the limited sections (up to 4), it was not feasible to delineate individual HVC areas. Instead, HVCs were divided into three subregions, AVC, MVC and LVC, based on their anatomical location relative to V1. The correspondence of these subregions to standard higher-order visual area nomenclature in the Allen Common Coordinate Framework is illustrated in Supplementary Fig. 1.

Third, to refine the borders between HVCs and other cortical areas, the spatial distribution of mCherry^+^ cells was used. As noted above, the spatial distribution of mCherry+ cells was relatively homogenous within subregions of HVCs, facilitating the delineation of HVA borders. Furthermore, borders between the MVC and retrosplenial cortex were refined using the distribution of supertype L2/3 IT RSP 1, which is selectively located in the retrosplenial cortex but not in the MVC.

After delineating the borders of V1, AVC, MVC, and LVC, regions of interest (ROI) within these cortical areas were manually defined using a customized MATLAB script for subsequent analyses of the molecular identity of neurons receiving transsynaptic mWGA-mCherry labeling. Because the dLGN and the LP are anatomically adjacent, viral injections of AAV mWGA-mCherry were restricted in volume to confine expression to the targeted nucleus and minimize spillover into the neighboring nucleus. As a result, viral expression did not span the entire dLGN or LP, and cortical regions innervated by unlabeled thalamic neurons within these thalamic nuclei lacked detectable mCherry signal. To ensure consistency in downstream analyses, in each cortical area, only the region exhibiting a spatially homogeneous mCherry signal was included, whereas the region within the same area exhibiting a substantially weaker or undetectable mCherry signal was discarded. In addition, a narrow band (∼20 µm) adjacent to the identified borders between cortical areas was excluded to minimize potential contamination from neighboring cortical areas.

### Fraction of mCherry^+^ neurons, GABAergic-to-glutamatergic ratio, and relative preference

For each animal, ROIs of the same cortical area were pooled across all applicable sections (up to 4). For each ROI in each animal, the fraction of mCherry^+^ neurons was computed by summing the posterior probabilities of mCherry positivity (as described in the section “Identify cells with mWGA-mCherry transfer”) across all neurons within each class, subclass, or supertype. Fractions of mCherry^+^ neurons from the same cortical area were then averaged across multiple animals within the same experimental group. Prior to statistical testing, fraction data were logit-transformed to stabilize variance and approximate normality. Comparisons of the overall fraction of mCherry^+^ neurons between two groups were assessed using Welch’s t-tests, whereas comparisons across multiple regions or conditions were evaluated using one-way ANOVA followed by Benjamini–Hochberg false discovery rate (FDR) correction for multiple testing. For subclass-level analyses, a two-way ANOVA was then applied to assess the effects of experimental condition, subclass type, and their interaction. To identify subclass-specific differences between conditions, pairwise comparisons were performed using Welch’s t-tests on the logit-transformed data, followed by FDR correction for multiple testing.

GABAergic-to-glutamatergic ratio was quantified as (total number of mCherry^+^ GABAergic neurons) / (total number of mCherry^+^ glutamatergic neurons) across all neurons or for neurons in each layer separately. Prior to statistical testing, GABAergic-to-glutamatergic ratios were log-transformed to stabilize variance. Differences in I/E ratios across regions were assessed using Welch’s t or one-way ANOVA followed by FDR correction for multiple testing.

Relative preference for each subclass or supertype was defined as their fraction of mCherry^+^ neurons in that group divided by the global fraction of mCherry^+^ neurons across the corresponding reference population. For example, the relative preference of each GABAergic subclass was calculated as the fraction of mCherry^+^ neurons within that subclass, divided by the fraction of mCherry^+^ neurons across all GABAergic neurons. Neurons belonging to minor subclasses that were excluded by filtering were not included in the calculation of the total mCherry⁺ fraction across all glutamatergic or GABAergic neurons. Prior to statistical testing, relative preference values were log_2_-transformed to achieve symmetry around zero (no enrichment or depletion) and approximate normality. To assess whether relative preference profiles differed between experimental groups, a two-way ANOVA was performed, with the interaction term indicating group-dependent shifts in cell-type targeting. For individual subclasses or supertypes, pairwise differences between conditions were evaluated with FDR correction for multiple comparisons. Within each group, enrichment or depletion relative to the reference population was assessed using one-sample t-tests against zero, also with FDR correction.

### Cell-type composition and gene expression similarity to V1

All mice with LP injections that enabled the delineation of HVA borders were included in this analysis (i.e., Emx1-Cre with LP injections and Calb2-Cre; n = 10).

To compute cell-type composition similarity, frequency vectors were constructed for V1 and each target region for each animal, and similarity was calculated from these vectors. Similarity was quantified using the Bray–Curtis metric, defined as 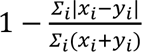, where *x_i_* and *y_i_* denote the frequencies of cell type i in the two regions, with values ranging from 0 (no shared composition) to 1 (identical composition). Analyses were performed at both subclass and supertype levels, and separately for glutamatergic and GABAergic populations.

To compute gene expression similarity, pseudo-bulk expression vectors were generated for each region in each animal by averaging gene expression across all cells within a given ROI. Pairwise similarity between regions for each animal was then quantified using two metrics:

The first metric was the Spearman’s rank correlation coefficient. To calculate it, genes were ranked by expression from lowest to highest for both V1 and the target HVA subregion. Spearman’s rank correlation coefficient was defined as 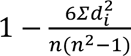, where *d_i_* is the difference between the two ranks for gene i and *d_i_* is the total number of genes. This metric reduces the influence of high-magnitude genes and is therefore robust to outliers.

The second metric was cosine similarity, defined as 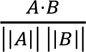, where *A* and *B* represent the pseudo-bulk gene expression vectors for V1 and the target HVA subregion. This metric captures the similarity in the pattern of gene expression independent of absolute magnitude.

Statistical significance for all similarity metrics (Bray–Curtis compositional similarity, Spearman correlation, and cosine similarity) was assessed using the same pipeline. Prior to hypothesis testing, similarity values were transformed to approximate normality: Spearman correlation coefficients were subjected to Fisher’s Z-transformation, whereas Bray–Curtis and cosine similarity values were logit-transformed. To compare similarity across HVA subregions, one-way ANOVA was applied to the transformed values, followed by post hoc pairwise comparisons with FDR correction for multiple testing.

## Data Availability

All the data that support the findings presented in this study are available from the corresponding author upon reasonable request.

## Code Availability

All Python and MATLAB scripts used in this manuscript will be shared via GitHub.

## Acknowledgement

We thank Li Wang for technical assistance and discussions, Xiaotian Wu for assistance with statistical analysis, Pooja Saraf, Lucy Pena, Jitao Hu for technical assistance, and current and past members of the Scanziani lab for discussions. Funding: National Institutes of Health grants R01 NS123912 (M.S.) and Howard Hughes Medical Institute (M.S.); Jane Coffin Childs Postdoctoral Fellowship (X.G.), NIH K99EY038331 (X.G.); NIH R01EY030138 (X.D.), R01 NS123912 (X.D.), U01NS136405 (X.D.), GRF-CFC3 (X.D.) and RPB-Stein Award (X.D.).

## Author Contributions

X.G., X.W., X.D., and M.S. designed the study. X.G., X.W., B.R., and Q.Y.W. conducted all experiments. X.G. analyzed the MERFISH data. X.W. analyzed the electrophysiological data. N.T. S.Z. and X.D. established the TransA-MERFISH protocol, including histology. S.Y.C. produced new AAV vectors in the study. X.G., X.W., X.D. and M.S. wrote the paper.

## These authors contributed equally: Xinxin Ge, Xing Wei

## Extended Data Figures 1-16

**Extended Data Figure 1.**
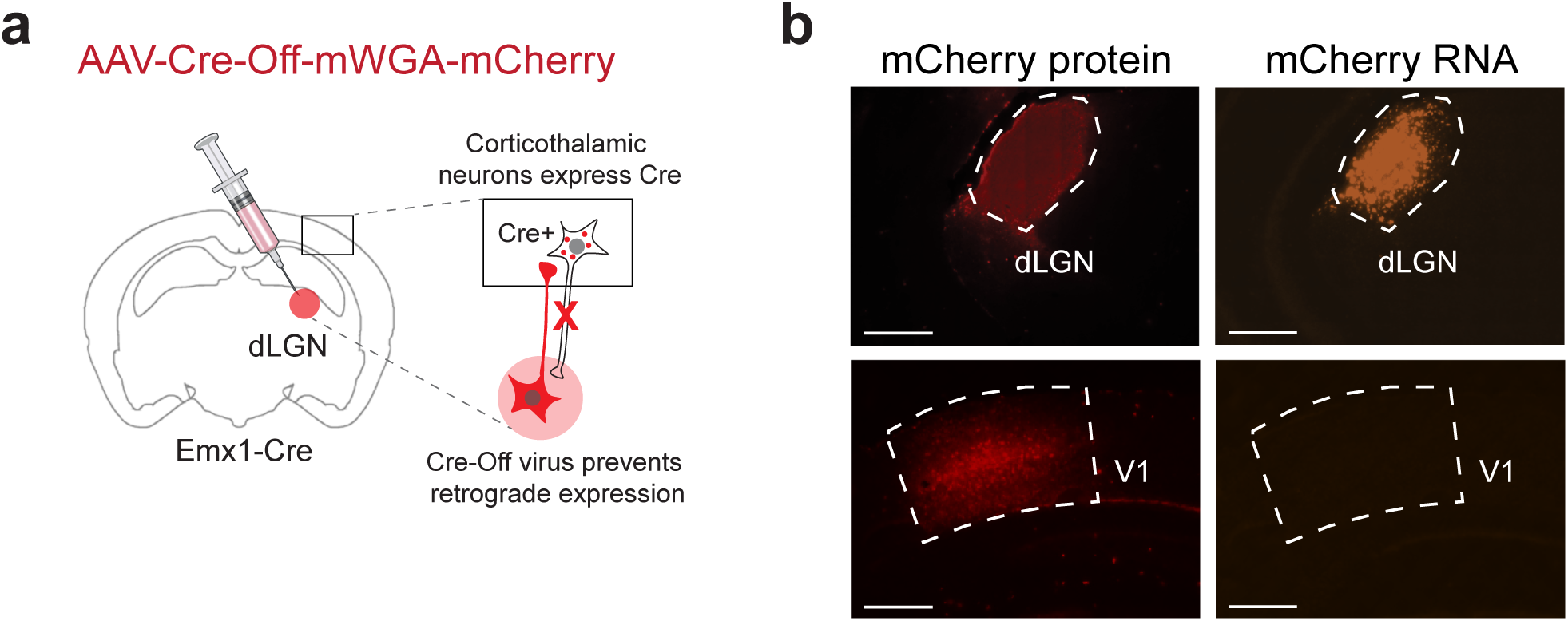
Cre-Off system prevents retrograde labelling. **a**, Schematic illustrating the Cre-Off approach (see Methods). **b**, MERFISH sections of the dLGN and V1 from the same animal. Top left, mWGA-mCherry protein staining in dLGN; top right, mWGA-mCherry RNA transcript signal in dLGN; bottom left, mWGA-mCherry protein staining in V1; bottom right, mWGA-mCherry RNA transcript signals in V1. Note the absence of mWGA-mCherry RNA signal in V1. Scale bars, 0.5mm.

**Extended Data Figure 2.**
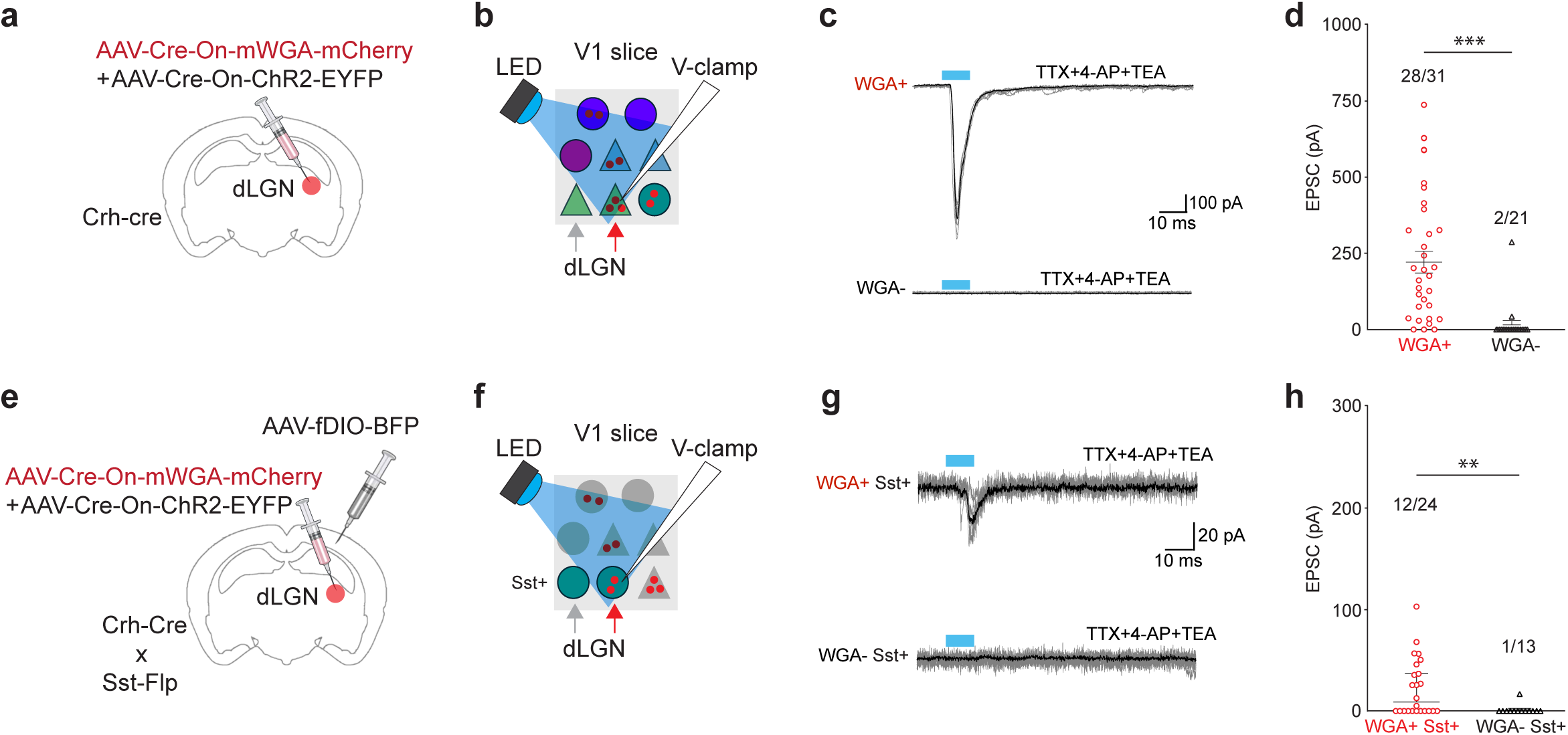
mWGA-mCherry transfer reveals mono-synaptic connectivity. **a**, Schematic illustrating the viral expression strategy (see Methods). **b**, Schematic of the recording configuration: whole cell recordings from mWGA-mCherry positive (and neighboring mWGA-mCherry negative) V1 neurons in a brain slice. **c**, Example recordings showing monosynaptic responses evoked by blue LED stimulation of dLGN axons. The mWGA-mCherry⁺ neuron (top) exhibits robust EPSCs, whereas a neighboring mWGA–mCherry- neuron (bottom) shows no detectable response. Recordings were performed in the presence of TTX, 4-AP, and TEA to isolate monosynaptic excitation. Black traces indicate the average of 8–10 sweeps (gray). Blue bars, 10ms blue LED pulses. **d**, Distribution of excitatory postsynaptic current (EPSC) amplitude recorded in mWGA-mCherry positive (red symbols) and negative (black symbols) V1 neurons. Note the low number of false positive (2/21) neurons. Neurons were pooled from n = 4 mice. Significance was determined using a two-sided Mann–Whitney U-test. **e-h,** same as **a-d**, but recordings were performed in Sst^+^ neurons using the viral expression strategy illustrated in **e**. Neurons were pooled from n = 6 mice. Bars represent a median ± 95% confidence interval across neurons. \**P* < 0.05, \*\**P* < 0.001, \*\*\**P* < 0.0001. Full details on statistical models and *P* values are provided in Supplementary Table 4.

**Extended Data Figure 3.**
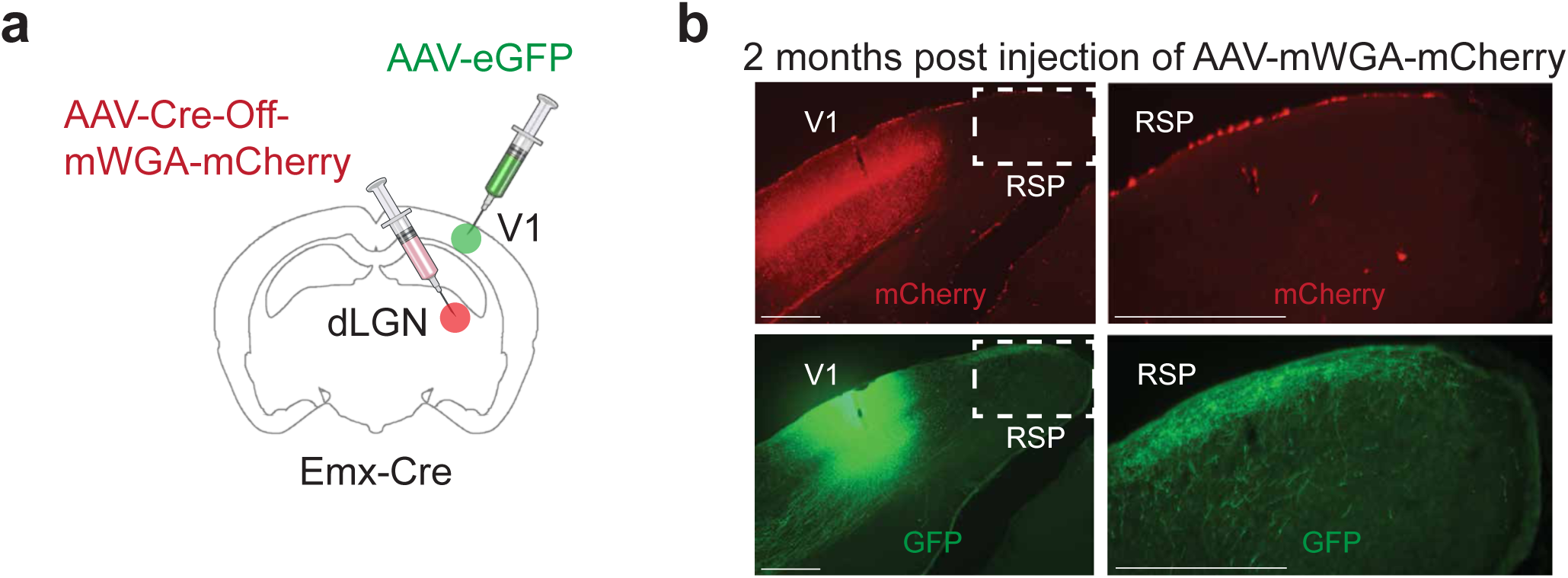
mWGA-mCherry anterograde transsynaptic tracing shows no secondary spread up to 2 months post injection. **a**, Schematic of the viral expression strategy (see Methods). **b**, Top left, mWGA-mCherry labeling is observed in V1 and not in the retrosplenial cortex (RSP) two months after AAV-mWGA-mCherry injection in the dLGN; top right, higher magnification of the RSP region outlined in top left. Bottom left, expression of eGFP in the same section, 2 weeks after AAV-eGFP injection in V1, to highlight the projection of V1 to RSP; bottom right, higher-magnification of the RSP region outlined in bottom left. Scale bars, 0.5mm.

**Extended Data Figure 4.**
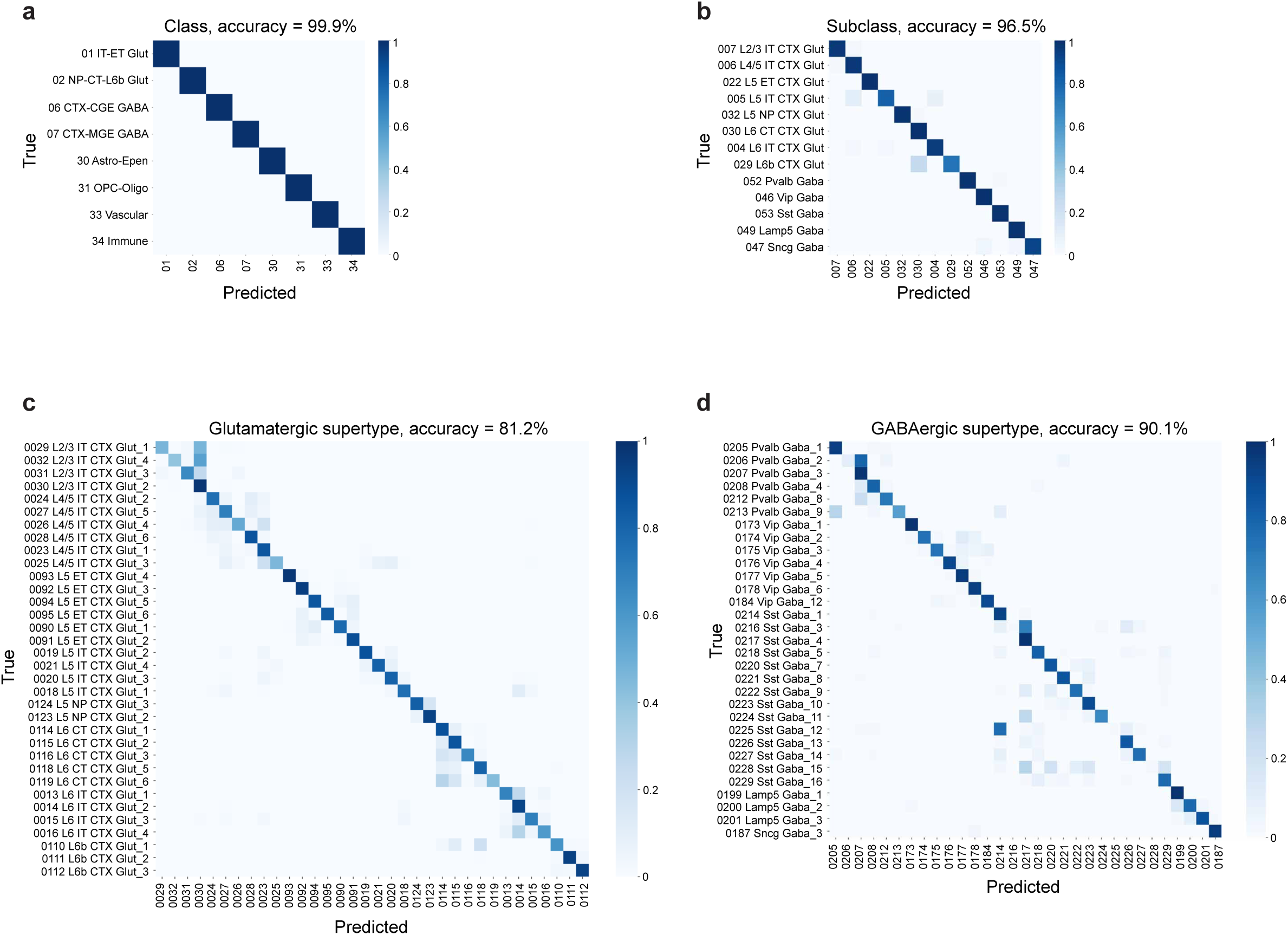
*In silico* validation of MERFISH panel performance in resolving visual cortical cell types. Row-normalized confusion matrices evaluating the predictive accuracy of a Random Forest classifier trained on the targeted 300-gene spatial panel at the class (**a**), subclass (**b**), and supertype levels for glutamatergic (**c**) and GABAergic (**d**) neurons (see Methods). For subclass and supertype analyses, only neuronal cell types were included. Consistent with downstream analyses, subclasses and supertypes with very low representation in the visual cortex were excluded (see Methods). Y-axis represents ground-truth cell identities derived from the single-cell RNAseq dataset (CCN20230722; Yao et al., 2023), x-axis represents the corresponding identities predicted by the classifier. Color intensity represents the fraction of true cells assigned to a given predicted label, with the dark diagonal indicating correct classifications. Glutamatergic subclasses and supertypes in **c** and **d** are ordered by cortical depth where applicable. Note that off-diagonal elements in **b**-**d** are largely confined to closely related or spatially adjacent sister populations.

**Extended Data Figure 5.**
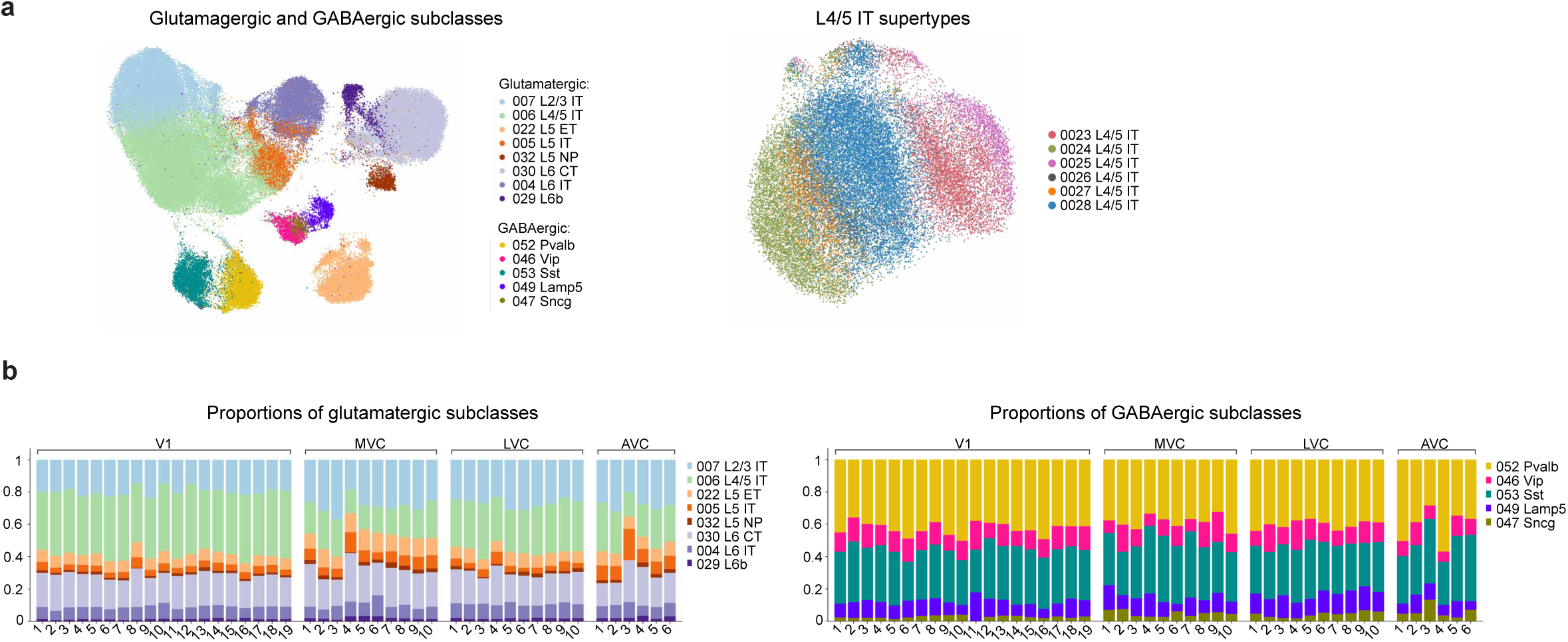
*In situ* validation of the MERFISH gene panel. **a.** UMAP based on the 300 gene panel used for MERFISH. Coloring denotes subclasses (left) and L4/5IT supertypes (right) based on the MapMyCell assignment. Note that neurons assigned to a given type (subclass or supertype) are close in UMAP space. **b.** Consistency of subclass distribution across animals. Left: Glutamatergic subclasses. Right GABAergic subclasses. Each column corresponds to a visual area in an animal.

**Extended Data Figure 6.**
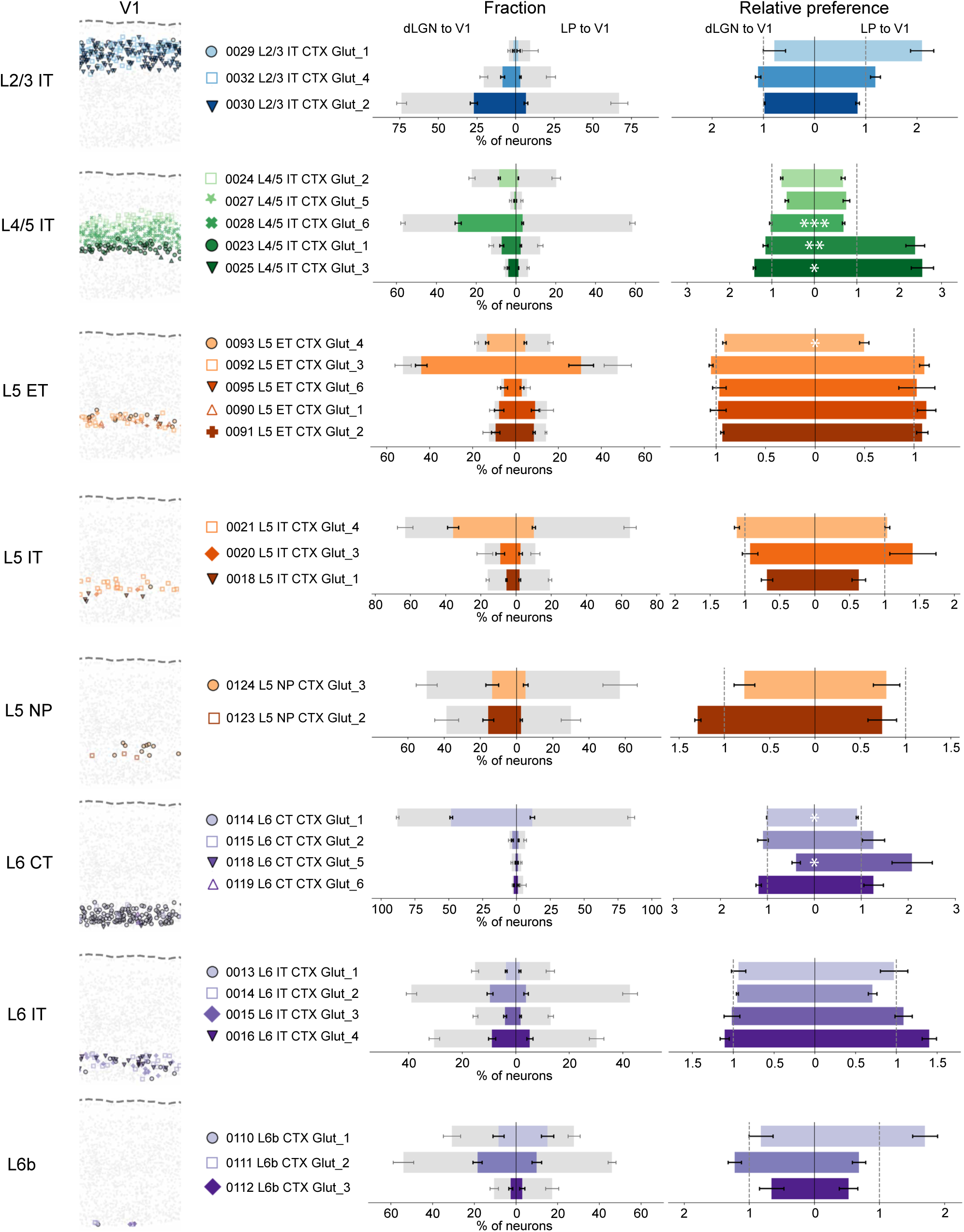
Relative preference of glutamatergic supertypes for dLGN and LP inputs to V1. Panels are organized by glutamatergic subclass from top to bottom. For each subclass, the spatial distribution (left), fraction of mCherry⁺ neurons (middle), and relative preference of individual supertypes (right). are shown. dLGN to V1: n=4, LP to V1: n=5. Relative preference of each supertype was calculated as the mCherry⁺ fraction in that supertype normalized to the mCherry⁺ fraction across all neurons in the same subclass. Supertypes are ordered by depth from the pia where applicable. Asterisks denote significant difference in relative preference between LP inputs and dLGN inputs for each glutamatergic supertype. Significance was assessed using two-sided Welch’s t-tests on log-transformed data, adjusted for multiple comparisons using FDR correction. Significance was only shown for subclasses with a significant interaction term in the two-way ANOVA after FDR correction. Bars represent mean ± s.e.m. across animals. \**P* < 0.05, \*\**P* < 0.001, \*\*\**P* < 0.0001. Full details on statistical models and *P* values are provided in Supplementary Table 4.

**Extended Data Figure 7.**
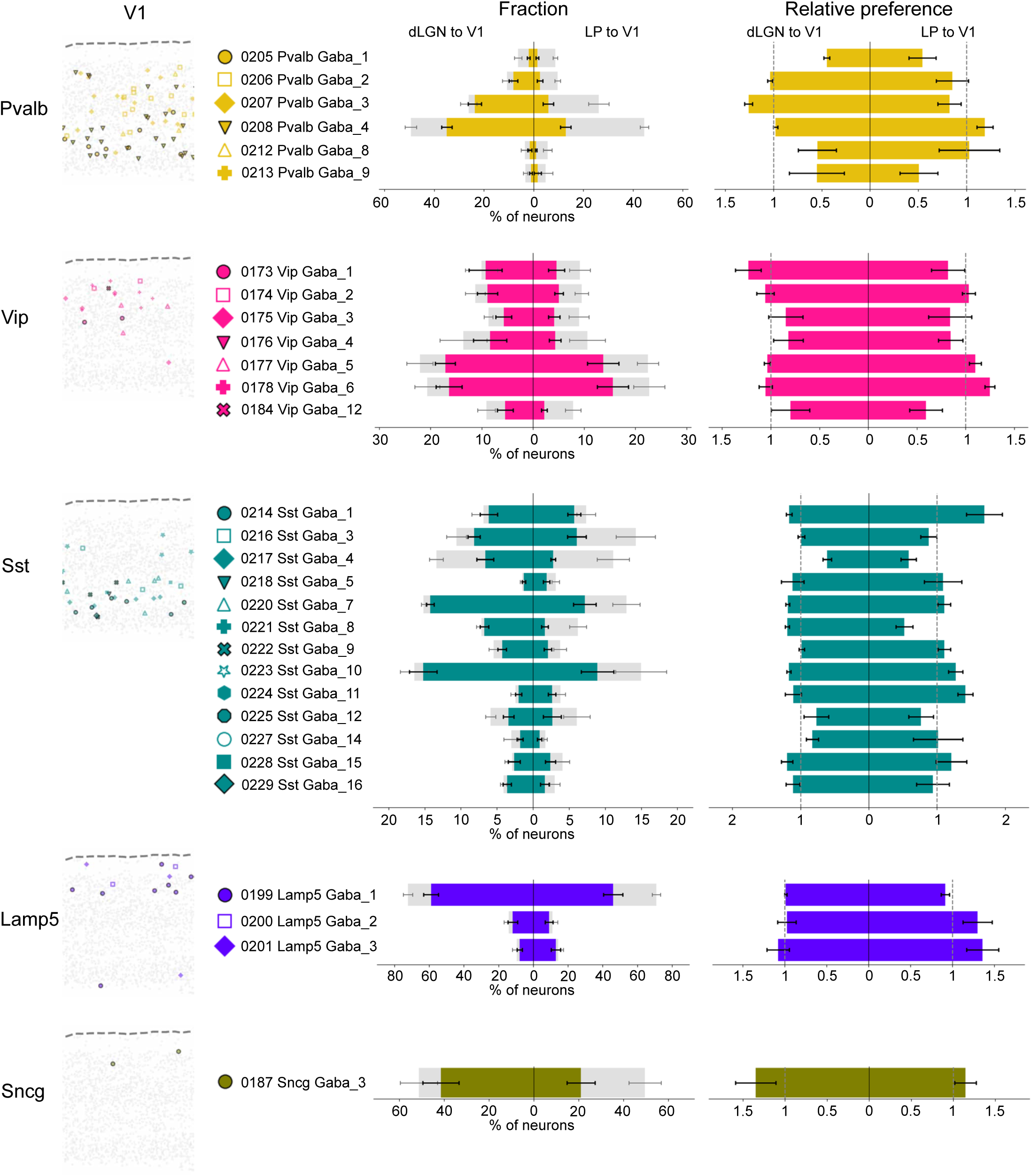
Relative preference of GABAergic supertypes for dLGN and LP inputs to V1. Panels are organized by GABAergic subclass from top to bottom. For each subclass, the spatial distribution (left), fraction of mCherry⁺ neurons (middle), and relative preference of individual supertypes (right) are shown. dLGN to V1: n = 4, LP to V1: n = 5. Relative preference of each supertype was calculated as the mCherry⁺ fraction in that supertype normalized to the mCherry⁺ fraction across all neurons in the same subclass. Significance was assessed using two-sided Welch’s t-tests on log-transformed data, adjusted for multiple comparisons using FDR correction. No subclass showed a significant interaction term in the two-way ANOVA after FDR correction. Bars represent mean ± s.e.m. across animals. Full details on statistical models and *P* values are provided in Supplementary Table 4. Note that despite significant differences in the relative preference of dLGN and LP for GABAergic subclasses in V1 (Fig. 2c) no significant differences were detected in the relative preferences of dLGN and LP for supertypes within these subclasses.

**Extended Data Figure 8.**
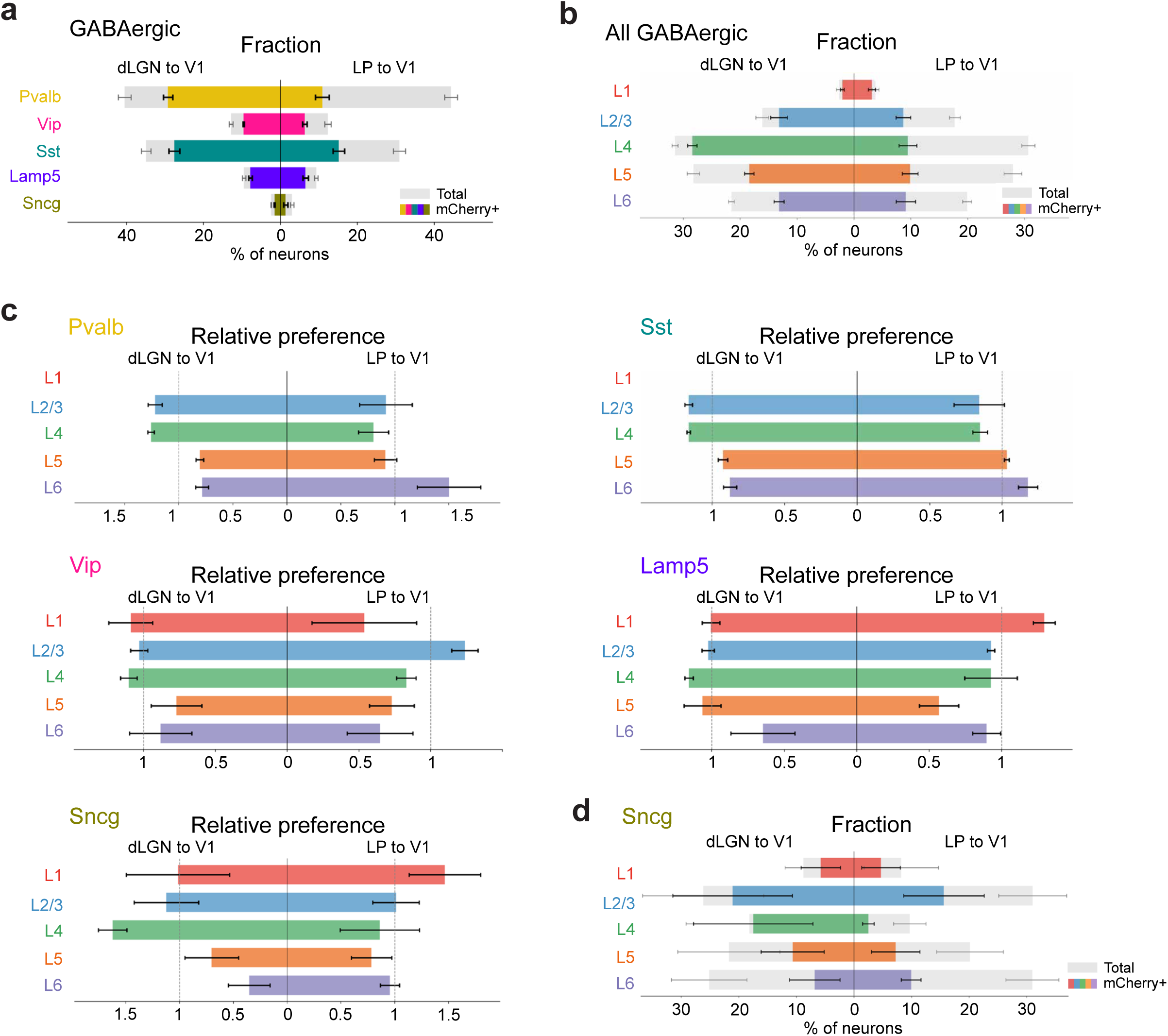
Fraction and relative preference of GABAergic neurons across cortical layers for dLGN and LP inputs to V1. **a,** Fractions of mCherry⁺ neurons across V1 GABAergic subclasses following dLGN (left) or LP (right) injections. **b,** Fractions of GABAergic mCherry⁺ neurons across cortical layers following dLGN (left) or LP (right) injections. **c**, Relative layer preferences of dLGN and LP inputs to V1 for each GABAergic subclass. **d**, Fractions of mCherry^+^ neurons of Sncg subclass across cortical layers in V1 following dLGN injections or LP injections. dLGN to V1: n = 4, LP to V1: n = 5. Bars represent mean ± s.e.m. across animals. \**P* < 0.05, \*\**P* < 0.001, \*\*\**P* < 0.0001. Statistical significance for mCherry⁺ fraction plots in **a, b, d** is not shown. Full details on statistical models and *P* values are provided in Supplementary Table 4.

**Extended Data Figure 9.**
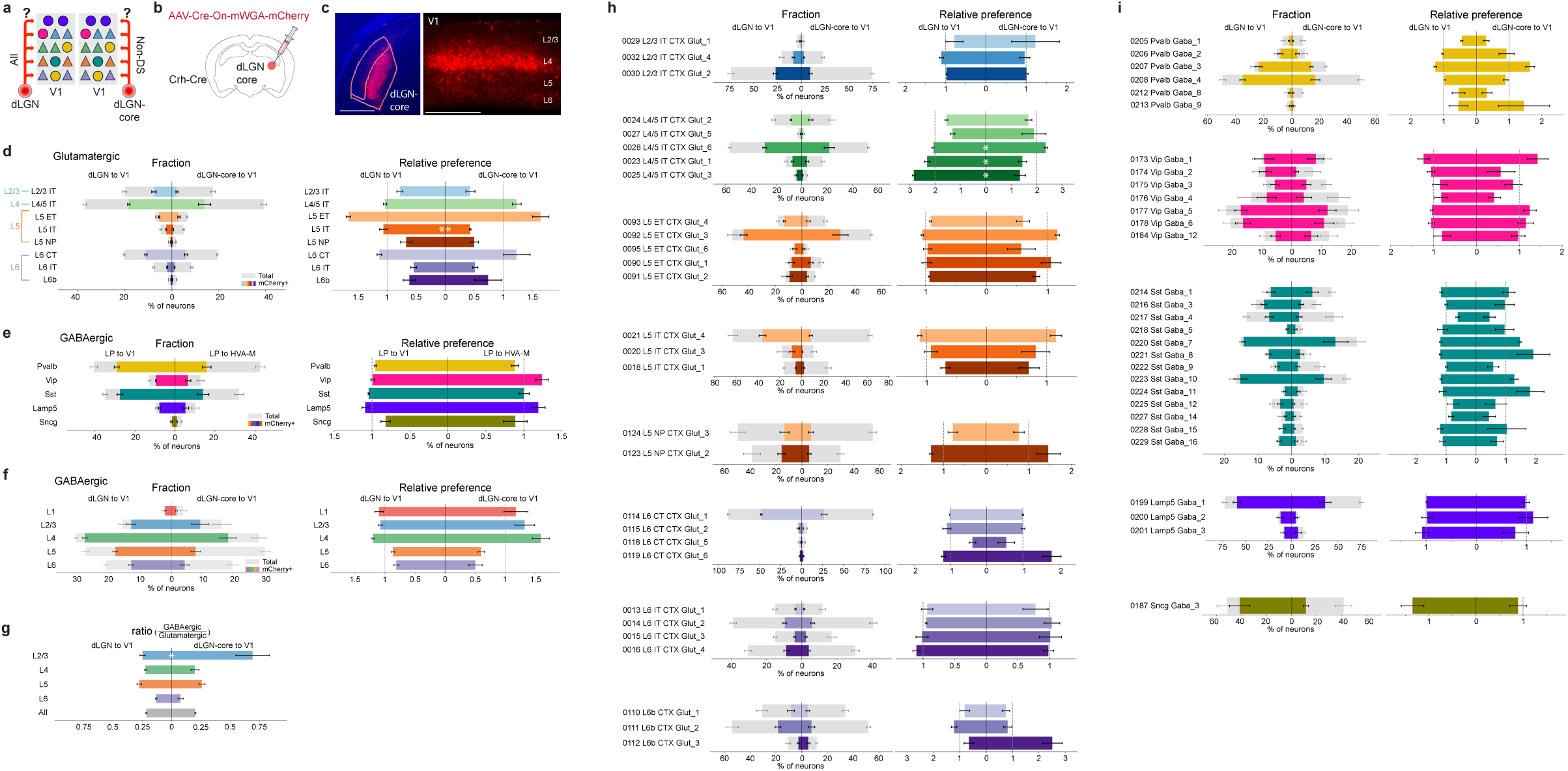
Patterns of V1 neurons targeted by Crh^+^ dLGN-core versus the overall dLGN input. **a**, Schematic illustrating the question: which molecularly defined neurons are targeted by the overall dLGN or Crh^+^ dLGN-core neurons? **b**, Schematic of the viral expression strategy. **c**, Representative images showing core-restricted mWGA-mCherry expression at the injection site in the thalamus (left) and anterogradely transferred mWGA-mCherry signal in V1 (right), three weeks after injection into the dLGN. **d, e,** Left, fractions of mCherry⁺ neurons across V1 glutamatergic (**d**) and GABAergic (**e**) subclasses comparing dLGN and Crh^+^ dLGN-core inputs to V1. Right, corresponding relative preferences for each glutamatergic (**d**) and GABAergic (**e**) subclass in V1. **f**, Fractions (left) and relative preferences (right) of GABAergic (all subclasses combined) neurons across cortical layers targeted by overall dLGN and Crh^+^ dLGN-core neurons. **g**, GABAergic-to-glutamatergic ratios of V1 neurons targeted by overall dLGN and Crh^+^ dLGN-core inputs, shown for each layer (colored) and across all layers (gray). **h, i**, Fractions (left) and relative preferences (right) of supertypes within each glutamatergic (**h**) and GABAergic (**i**) subclass comparing overall dLGN and Crh^+^ dLGN-core inputs. Panels are organized by subclass from top to bottom. Overall dLGN to V1: n = 4, Crh^+^ dLGN-core to V1: n = 5. Significance shown only for relative preference and GABAergic-to-glutamatergic ratio, determined using two-sided Welch’s t-tests on log-transformed data, adjusted for multiple comparisons using FDR correction. For **h** and **i**, significance was only shown for subclasses with a significant interaction term in the two-way ANOVA after FDR correction. Bars represent mean ± s.e.m. across animals. \**P* < 0.05, \*\**P* < 0.001, \*\*\**P* < 0.0001. Full details on statistical models and *P* values are provided in Supplementary Table 4.

**Extended Data Figure 10.**
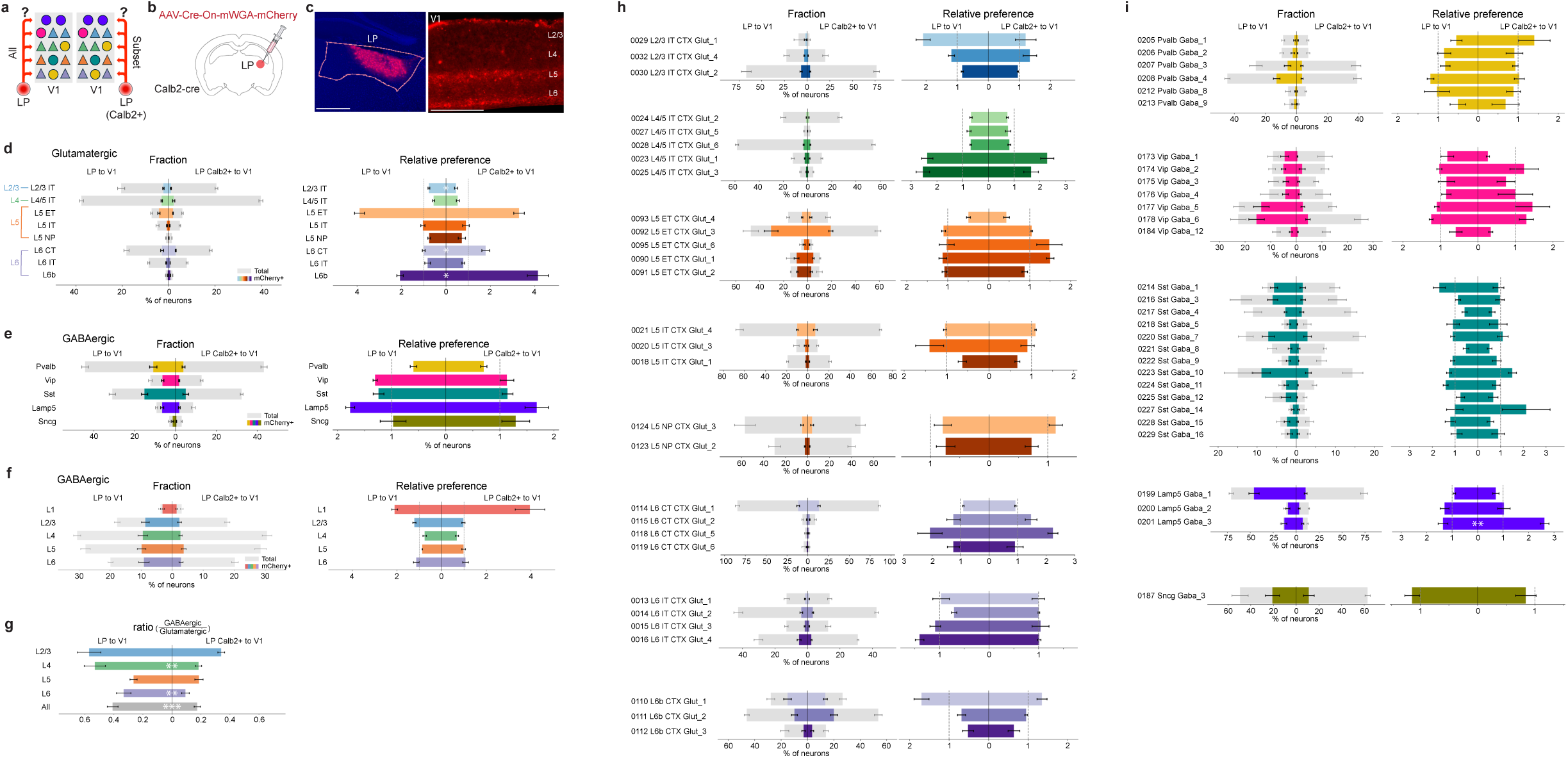
Patterns of V1 neurons targeted by Calb2⁺ LP versus the overall LP input. **a**, Schematic illustrating the question: which molecularly defined neurons in V1 are targeted by the overall LP or Calb2^+^ LP neurons? **b**, Schematic of the viral expression strategy. **c**, Representative images showing mWGA-mCherry expression at the injection site in the thalamus and anterogradely transferred mWGA-mCherry signal in V1, three weeks after LP injection. **d, e,** Left, fractions of mCherry⁺ neurons across V1 glutamatergic (**d**) and GABAergic (**e**) subclasses comparing LP and Calb2^+^ LP inputs to V1. Right, corresponding relative preferences for each glutamatergic (**d**) and GABAergic (**e**) subclass in V1. **f**, Fractions (left) and relative preferences (right) of GABAergic neurons (all subclasses combined) across cortical layers targeted by overall LP and Calb2^+^ LP neurons. **g**, GABAergic-to-glutamatergic ratios of V1 neurons targeted by overall LP and Calb2^+^ LP inputs, shown for each layer (colored) and across all layers (gray). **h, i**, Fractions (left) and relative preferences (right) of supertypes within each glutamatergic (**h**) and GABAergic (**i**) subclass comparing overall LP and Calb2^+^ LP inputs. Panels are organized by subclass from top to bottom. Overall LP to V1: n = 5, Calb2^+^ LP to V1: n = 5. Significance shown only for relative preference and GABAergic-to-glutamatergic ratio, determined using two-sided Welch’s t-tests on log-transformed data, adjusted for multiple comparisons using FDR correction. For **h** and **i**, significance was only shown for subclasses with a significant interaction term in the two-way ANOVA after FDR correction. Bars represent mean ± s.e.m. across animals. \**P* < 0.05, \*\**P* < 0.001, \*\*\**P* < 0.0001. Full details on statistical models and *P* values are provided in Supplementary Table 4.

**Extended Data Figure 11.**
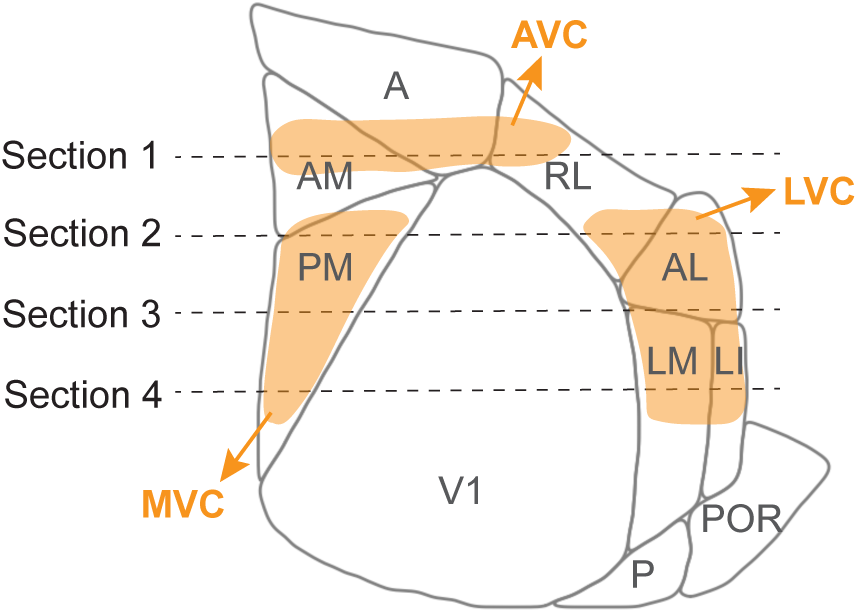
Mapping MVC, AVC, and LVC to the Allen Common Coordinate Framework. Correspondence of MVC, AVC, and LVC (in orange) to standard higher-order visual area nomenclature in the Allen Common Coordinate Framework (CCFv3). Dashed lines indicate the estimated anterior–posterior positions of the collected coronal sections.

**Extended Data Figure 12.**
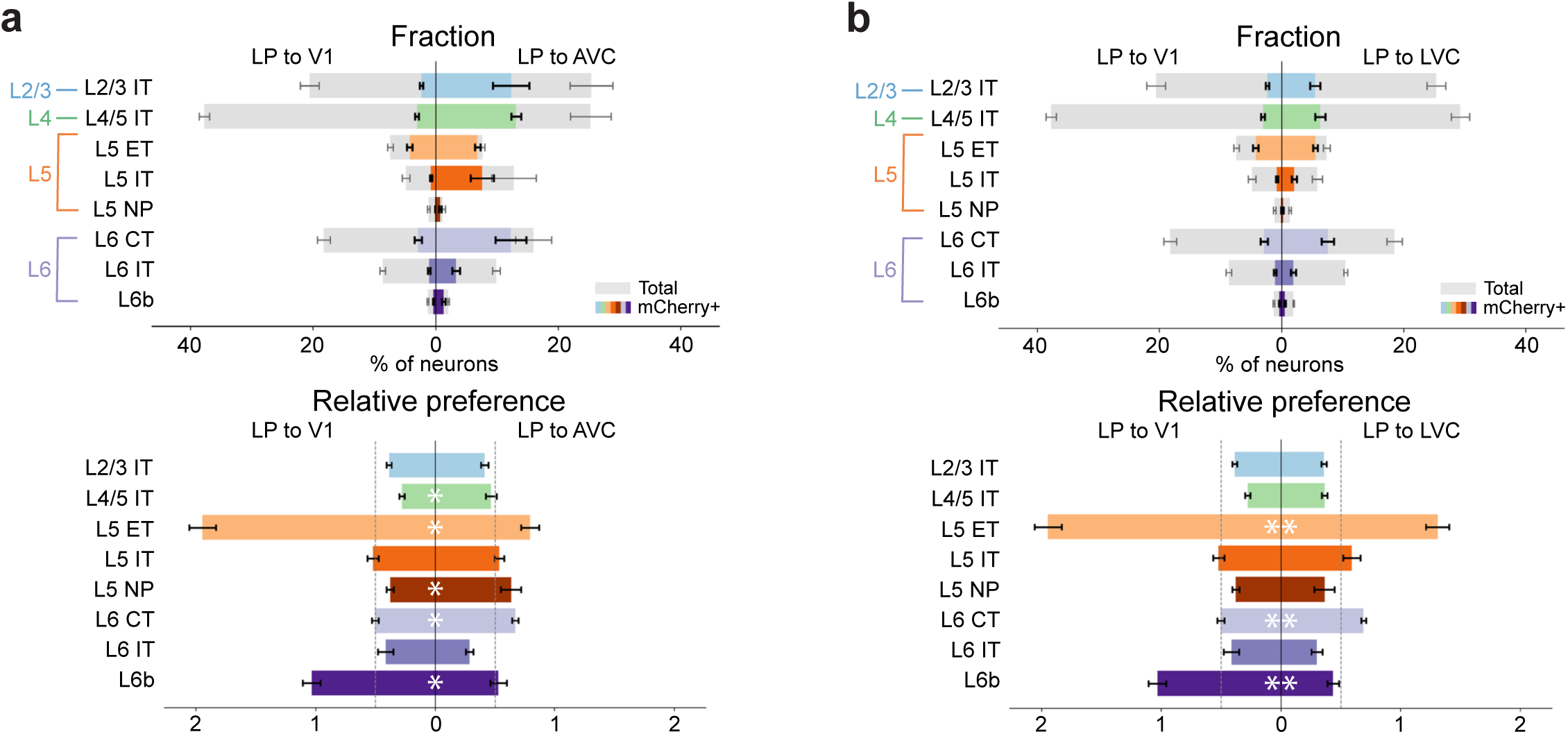
Fractions and relative preferences of glutamatergic subclasses targeted by LP inputs to V1 compared to LP input to AVC and LVC. **a, b,** Top, fractions of mCherry^+^ neurons across glutamatergic subclasses in V1 and AVC (**a**) or LVC (**b**) following LP injections. Bottom, corresponding relative preference values for each glutamatergic subclass comparing LP inputs to V1 (n = 5) and AVC (n = 3) (**a**) or LVC (n = 5) (**b**). LP inputs to V1, AVC and LVC were measured in the same animals, with two animals lacking AVC data. Significance was determined using two-sided Welch’s t-tests on log-transformed data, adjusted for multiple comparisons using FDR correction. Significance shown only for relative preference. Bars represent mean ± s.e.m. across animals. \**P* < 0.05, \*\**P* < 0.001, \*\*\**P* < 0.0001. Full details on statistical models and *P* values are provided in Supplementary Table 4.

**Extended Data Figure 13.**
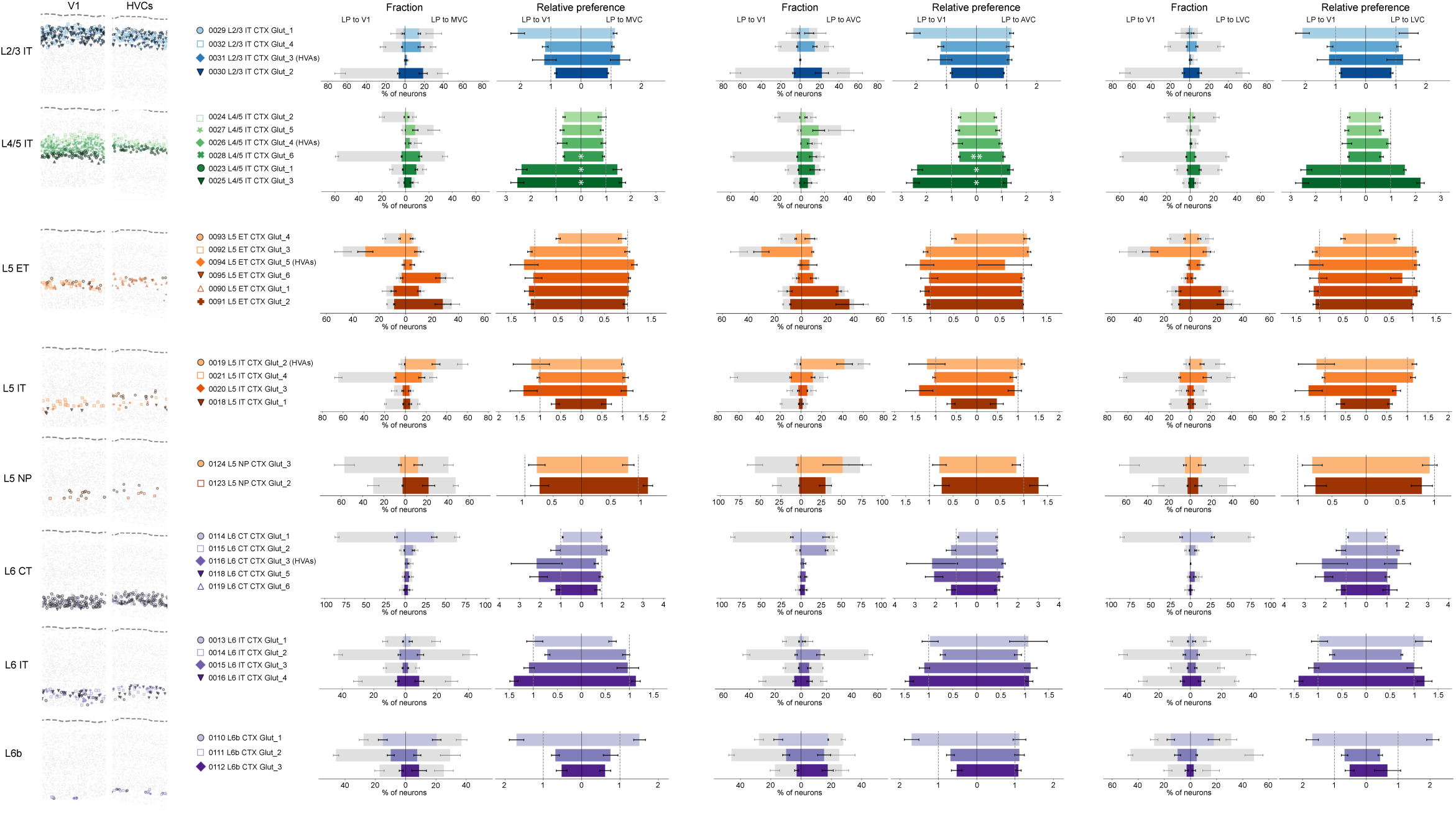
Fractions and relative preference of glutamatergic supertypes targeted by LP inputs to V1, compared to LP input to MVC, AVC and LVC. Panels are organized by glutamatergic subclass from top to bottom. For each subclass, the spatial distribution of individual supertypes is shown at left, followed by the fraction of mCherry⁺ neurons and the relative preference for each supertype in LP to V1 (n = 5) versus LP to MVC (n = 5), LP to AVC (n = 3) and LP to LVC (n = 5). LP inputs to V1, LVC, AVC and MVC were measured in the same animals, with two animals lacking AVC data. Significance was assessed using two-sided Welch’s t-tests on log-transformed data, adjusted for multiple comparisons using FDR correction. Significance is shown only for relative preference, and only for subclasses with a significant interaction term in the two-way ANOVA after FDR correction. Bars represent mean ± s.e.m. across animals. \**P* < 0.05, \*\**P* < 0.001, \*\*\**P* < 0.0001. Full details on statistical models and *P* values are provided in Supplementary Table 4. Note that although significance is not shown for L5 ET supertypes because the two-way ANOVA interaction was not significant after FDR correction, supertype 0093, identified in prior analyses, nevertheless differed significantly between LP to V1 and LP to MVC or LP to AVC, but not LP to LVC.

**Extended Data Figure 14.**
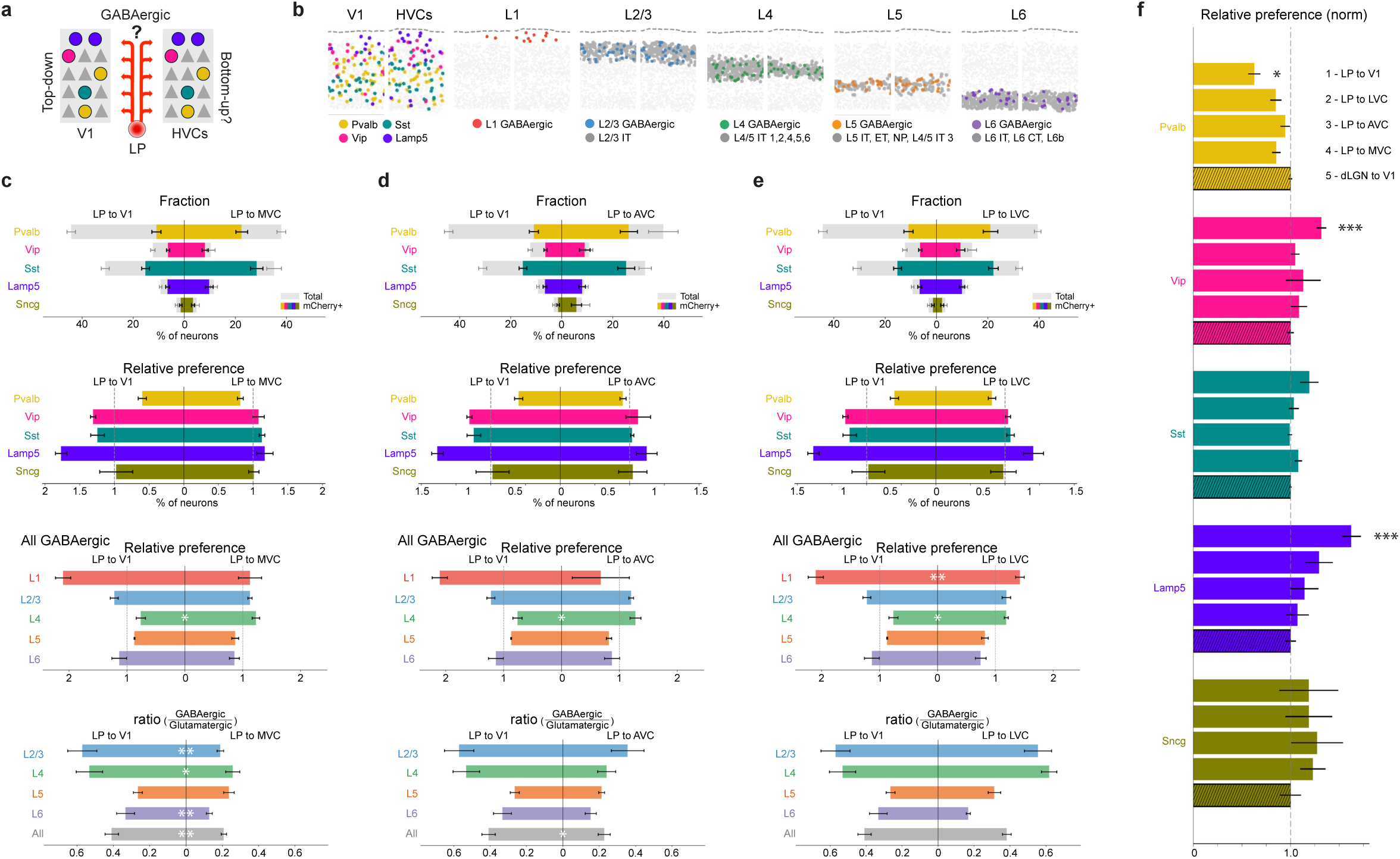
GABAergic subclasses in V1 and HVCs targeted by LP inputs. **a**, Schematic illustrating the question: which molecularly defined GABAergic neurons in V1 and in HVCs are targeted by LP input? **b**, Spatial organization of GABAergic neurons in example V1 and HVC sections. Left, spatial distribution of GABAergic subclasses; colored dots indicate neurons belonging to the subclasses listed below, and all other cells are shown in light gray. Right, spatial distribution of GABAergic neurons assigned to cortical layers (left to right, L1, L2/3, L4, L5 and L6). Colored dots indicate GABAergic neurons assigned to each layer, and dark gray dots indicate glutamatergic anchor cells used for proximity-based layer assignment. The subclass and supertype identities of the glutamatergic anchor cells used for each layer are indicated below. **c-e**, Top to bottom, comparisons of LP inputs to V1 versus LP inputs to MVC (c), AVC (d), and LVC (e), showing mCherry⁺ fractions in each GABAergic subclass, relative preference for each GABAergic subclass, relative preference of GABAergic neurons (all subclasses combined) across cortical layers, and GABAergic-to-glutamatergic ratios for each layer (colored) and across all layers (gray). Significance was determined using two-sided Welch’s t-tests on log-transformed data, adjusted for multiple comparisons using FDR correction. Significance for mCherry⁺ fraction plots is not shown. **f**, Normalized relative preference for GABAergic subclasses, comparing LP inputs to V1, LVC, AVC and MVC, and dLGN input to V1. LP inputs to V1, LVC and MVC were measured in the same animals, with two animals lacking AVC data. Relative preference values for each subclass were normalized to those of dLGN input to V1. Differences between LP inputs to V1, LVC, AVC and MVC and dLGN input to V1 were assessed using two-sided Welch’s t-tests on log-transformed data, with FDR correction for multiple comparisons. LP to V1: n = 5, LP to LVC: n = 5, LP s to AVC: n = 3, LP to LVC: n = 5, dLGN to V1: n = 4. LP inputs to V1, LVC, AVC and MVC were measured in the same animals, with two animals lacking AVC data. Bars represent mean ± s.e.m. across animals. \**P* < 0.05, \*\**P* < 0.001, \*\*\**P* < 0.0001. Full details on statistical models and *P* values are provided in Supplementary Table 4. Note that the LP innervation patterns that differ the most from the dLGN to V1 innervation pattern are LP to V1, consistent with the LP innervation of glutamatergic subclasses (Fig. 3e).

**Extended Data Figure 15.**
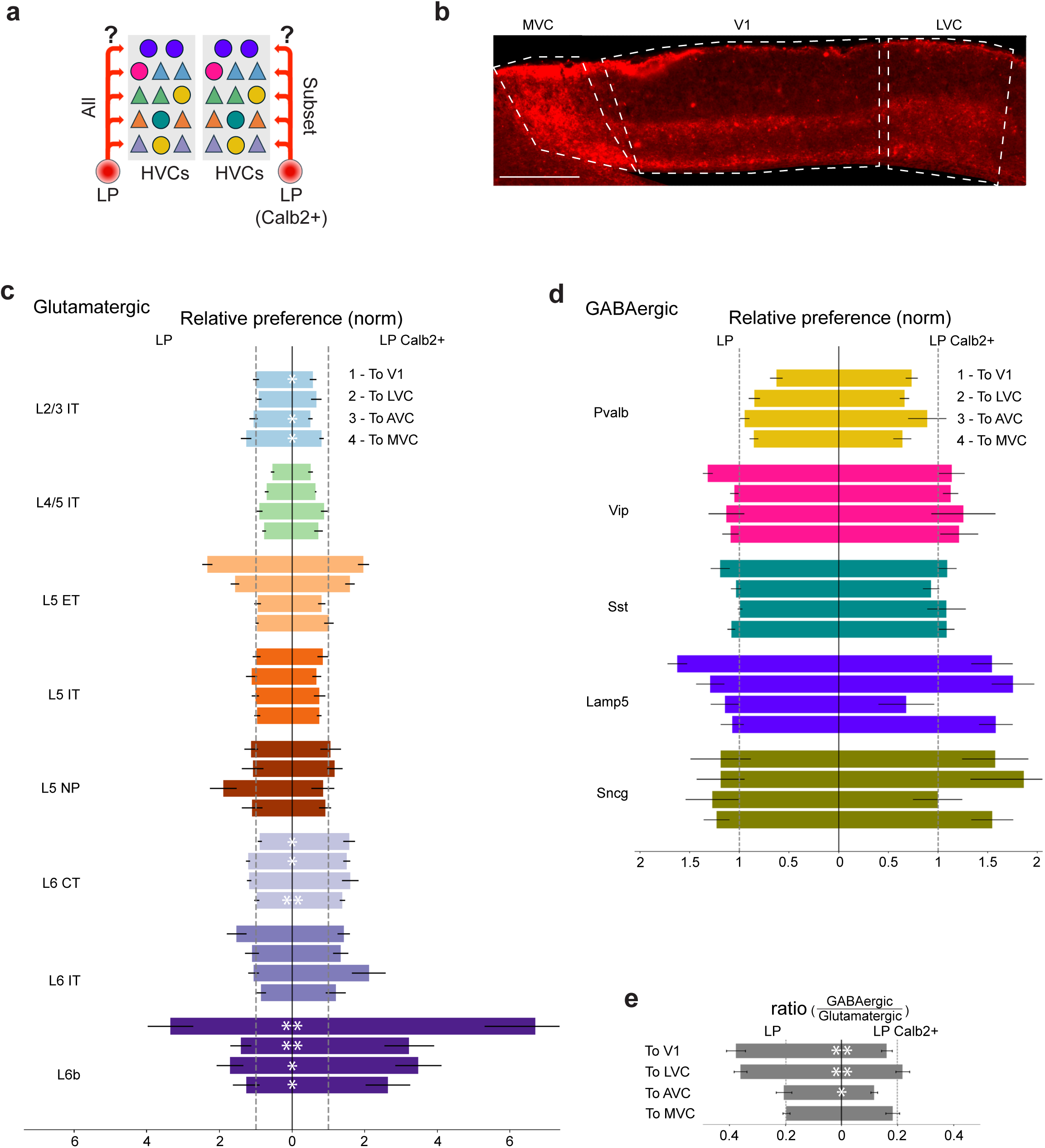
Patterns of HVC neurons targeted by Calb2⁺ LP versus overall LP input. **a**, Schematic illustrating the question: which molecularly defined neurons in HVCs are targeted by the overall LP or Calb2^+^ LP neurons? **b**, Representative images showing anterogradely transferred mWGA-mCherry signal in V1 and HVCs, three weeks after LP injection. **c, d**, Normalized relative preference for glutamatergic (**c**) and GABAergic (**d**) subclasses, comparing overall LP input (left) versus Calb2^+^ LP input (right) to V1, LVC, AVC and MVC. Overall LP or Calb2^+^ LP inputs to V1, LVC, AVC and MVC were measured in the same animals, with two animals of each group lacking AVC data. Relative preference values for each subclass were normalized to those of dLGN input to V1, indicated by the gray dashed lines. **e**, GABAergic-to-glutamatergic ratios of V1, LVC, AVC and MVC neurons targeted by overall LP and Calb2^+^ LP inputs. N = 5 for V1, MVC and LVC, and 3 for AVC, for both overall LP and Calb2^+^ LP inputs. Within each dataset, measurements for V1, MVC, AVC and LVC were obtained from the same animals, with AVC data unavailable for two animals. Significance was determined using two-sided Welch’s t-tests on log-transformed data, adjusted for multiple comparisons using FDR correction. Bars represent mean ± s.e.m. across animals. \**P* < 0.05, \*\**P* < 0.001, \*\*\**P* < 0.0001. Full details on statistical models and *P* values are provided in Supplementary Table 4.

**Extended Data Figure 16.**
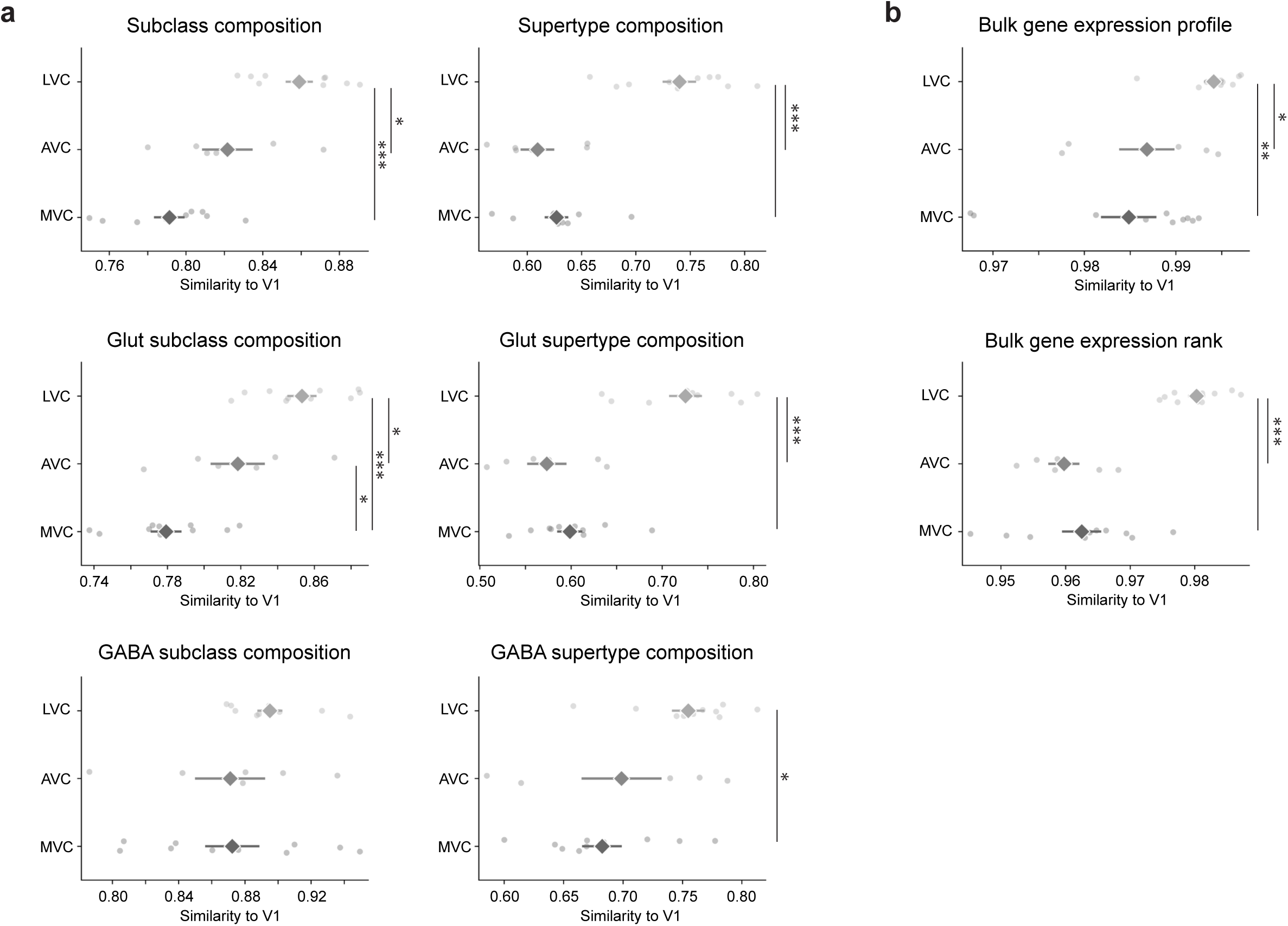
HVCs similarity to V1. **a**, Left, from top to bottom: subclass, glutamatergic subclass, and GABAergic subclass compositional similarity to V1, comparing LVC, AVC, and MVC, quantified using the Bray–Curtis metric (see Methods). Right, from top to bottom: supertype, glutamatergic supertype, and GABAergic supertype compositional similarity to V1. **b**, Top, cosine similarity of bulk gene expression to V1, comparing LVC, AVC, and MVC. Bottom, rank-based similarity of bulk gene expression to V1, quantified using Spearman’s rank correlation coefficient (see Methods). LVC: n = 10, AVC: n = 6, MVC: n = 10. LVC, AVC and MVC were measured in the same animals, with AVC data unavailable for four animals. Significance was determined using standard two-sided t-tests on logit-transformed (for Bray–Curtis and cosine similarity values) or Fisher’s Z-transformed data (for Spearman correlation coefficients), with FDR correction for multiple comparisons. Bars represent mean ± s.e.m. across animals. \**P* < 0.05, \*\**P* < 0.001, \*\*\**P* < 0.0001. Full details on statistical models and *P* values are provided in Supplementary Table 4.

